# Coronavirus hemagglutinin-esterase and spike proteins co-evolve for functional balance and optimal virion avidity

**DOI:** 10.1101/2020.04.03.003699

**Authors:** Yifei Lang, Wentao Li, Zeshi Li, Danielle Koerhuis, Arthur C.S. van den Burg, Erik Rozemuller, Berend-Jan Bosch, Frank J.M. van Kuppeveld, Geert-Jan P.H. Boons, Eric G. Huizinga, Hilde M. van der Schaar, Raoul J. de Groot

## Abstract

Human coronaviruses OC43 and HKU1 are respiratory pathogen of zoonotic origin that have gained worldwide distribution. OC43 apparently emerged from a bovine coronavirus (BCoV) spill-over. All three viruses attach to 9-*O*-acetylated sialoglycans via spike protein S with hemagglutinin-esterase HE acting as a receptor-destroying enzyme. In BCoV, an HE lectin domain promotes esterase activity towards clustered substrates. OC43 and HKU1, however, lost HE lectin function as an adaptation to humans. Replaying OC43 evolution, we knocked-out BCoV HE lectin function and performed forced evolution-population dynamics analysis. Loss of HE receptor-binding selected for second-site mutations in S, decreasing S binding affinity by orders of magnitude. Irreversible HE mutations selected for cooperativity in virus swarms with low-affinity S minority variants sustaining propagation of high-affinity majority phenotypes. Salvageable HE mutations induced successive second-site substitutions in both S and HE. Apparently, S and HE are functionally interdependent and co-evolve to optimize the balance between attachment and release. This mechanism of glycan-based receptor usage, entailing a concerted, fine-tuned activity of two envelope protein species, is unique among CoVs, but reminiscent of that of influenza A viruses (IAVs). Apparently, general principles fundamental to virion-sialoglycan interactions prompted convergent evolution of two important groups of human and animal pathogens.

## INTRODUCTION

The subfamily *Orthocoronavirinae* comprises a group of enveloped positive-strand RNA viruses of clinical and veterinary significance. Adding to the socio-economic impact of coronaviruses (CoVs) already extant in humans and livestock, the emergence of ‘new’ CoVs through cross species transmission poses an ever-looming threat to public health, animal health, and food production. Seven coronaviruses are known to infect humans, but not all of them have become established. The introduction of SARS CoV in 2002 from horseshoe bats with masked palm civets as incidental transient hosts, was rapidly contained through quarantine measures^1^. MERS CoV, natural to dromedary camels, causes a classical zoonotic infection with limited human-to-human spread^2^. December 2019, a member of the species *Severe acute respiratory syndrome related coronavirus* (SARS-CoV), called SARS-CoV-2 and 79.5% identical to the 2002 SARS CoV variant, emerged in Wuhan, China^3,4^ to progress to full scale pandemicity. Chances are, SARS-CoV-2 will eventually become established in the human population.

Four other respiratory coronaviruses of zoonotic origin already did succeed in becoming true human viruses with world-wide distribution^5–7^. Among them are HKU1 and OC43 (subgenus *Embecovirus*, genus *Betacoronavirus*), related yet distinct viruses that arose from different zoonotic progenitors and entered the human population independently. OC43 is far more related to bovine coronavirus (BCoV), its presumptive ancestor, with early isolates sharing 97% genome identity^8,9^. Together with viruses of swine, canines, equines and lagomorphs, OC43 and BCoV are considered host range variants of the virus species *Betacoronavirus-1* (collectively referred to as β1CoVs throughout)^7^. OC43 apparently emerged 70 to 130 years ago from a single cross species transmission event that gave rise to a human-only virus^8–10^. Like OC43, other β1CoVs also exhibit host specificity^8,11^. While these observations attest to the host promiscuity and zoonotic potential of embecoviruses and β1CoVs in particular, they are also indicative for the existence of host barriers, the breaching of which selects for adaptive mutations that result in host specialization and, ultimately, virus speciation. Conceivably, comparative studies of BCoV and OC43 may identify factors that promote or restrict cross species transmission of CoVs and thus further our understanding of the requirements for colonization of the human host.

Embecoviruses, OC43 and BCoV included, differ from other CoVs in that they encode two types of surface projections. Homotrimeric ‘peplomers’ comprised of spike protein S and extending 20 nm from the viral membrane, mediate virion attachment to entry receptors and membrane fusion^12^. Interspersed are stubby 8-nm homodimeric projections comprised of the hemagglutinin-esterase (HE)^13–15^, a dual function protein typically encompassing a receptor-binding lectin domain specific for *O*-acetylated sialic acid (*O*-Ac-Sia) and a receptor destroying sialate-*O*-acetylesterase domain^16–20^. The HE lectin domain contributes to virion attachment, but at the same time enhances sialate-*O*-acetylesterase activity towards clustered sialoglycotopes^11^.

Some embecoviruses, like mouse hepatitis virus (MHV) and related CoVs in rodents, attach to 4- or 9- *O*-acetylated sialosides (4- or 9-*O*-Ac-Sias) via HE^21–25^ and to a proteinaceous entry receptor via S^26,27^. Others, animal β1CoVs included, bind to 9-*O*-Ac-Sias via HE^28^ but, remarkably, also via S^29,30^ or, in the case of human coronaviruses OC43 and HKU1, even exclusively via S^11,31,32^.

Structure function analyses of HE and S proteins have yielded a wealth of data on ligand binding, substrate selection and protein-glycan interactions. The receptor binding sites (RBSs) of CoV HE lectin domains and those in related proteins of toro- and influenza C/D viruses ^22,33–35^ differ in sequence and structure yet conform to a common architectural design with a deep hydrophobic pocket (‘P1’) to accommodate the crucial sialate-*O*-acetyl moiety, and an adjacent pocket or depression (‘P2’) to accept the 5-*N*-acyl group^17,21,22,33^. Characteristically, P1 and P2 are separated by an aromatic side chain and binding of the ligand is stabilized further through electrostatic protein-glycan interactions typically involving distinctive Sia functions such as the Sia glycerol side chain, the 5-*N*-Acyl and/or the Sia carboxylate. The RBS for 9-*O*-Ac-Sia in the S proteins of BCoV and OC43, identified by comparative structural analysis^32^ and confirmed by the cryo-EM holostructure of OC43 S^36^, conforms to this blueprint. Moreover, this site is structurally and functionally conserved in HKU1^32^.

Much less is known about the functional relationship between S and HE, and the role of HE in particular remains poorly understood. In MHV, HE expression is dispensable for replication and rapidly lost during cell culture propagation^15^. Conversely, in β1CoVs, HE seems critical for infection. In OC43, loss of HE-associated acetyl esterase activity abrogates the production of infectious virus and virus dissemination in cell culture^37^. Moreover, acetyl-esterase inhibitors impede BCoV replication^19^, and antibodies against HE neutralize the virus *in vitro* and *in vivo*^38–40^. Still, even among β1CoVs there are differences in HE function apparently correlating with host specificity. Whereas HE lectin activity is strictly maintained in BCoV^28^, OC43 lost this function through progressive accumulation of mutations in the HE RBS, apparently as an adaptation to replication in the human respiratory tract^11^. Nevertheless, isolates of either virus propagate in cultured cells. To better understand the consequences of loss of HE lectin function as it occurred during OC43 and also HKU1 evolution, we took a reverse genetics/forced evolution approach with BCoV as a model. The findings reveal that HE and S are functionally interdependent and that the acquisition of HE by a proto-embecovirus allowed its β1CoV descendants to adopt strategies for reversible virion-sialoglycan attachment, remarkably similar to those of influenza A viruses.

## RESULTS

### Disruption of HE lectin function selects for mutations in S

To study the role and importance of HE in β1CoV propagation, we developed a reverse genetics system for BCoV strain Mebus (**sFig. 1A**) based on targeted RNA recombination^41,42^. Recombinant ‘wildtype’ BCoVs (rBCoV) with parental type HE and S, but with accessory ORF4a replaced by the *Renilla* luciferase gene (rBCoV-Rluc), were readily generated upon seeding acceptor-virus-infected, donor RNA-transfected mouse LR7 cells onto monolayers of feeder HRT18 cells. rBCoV-Rluc arose and within seven days grew to final titers routinely obtained for wildtype BCoV (_∼_10^8^ TCID50/ml).

**Fig. 1.**
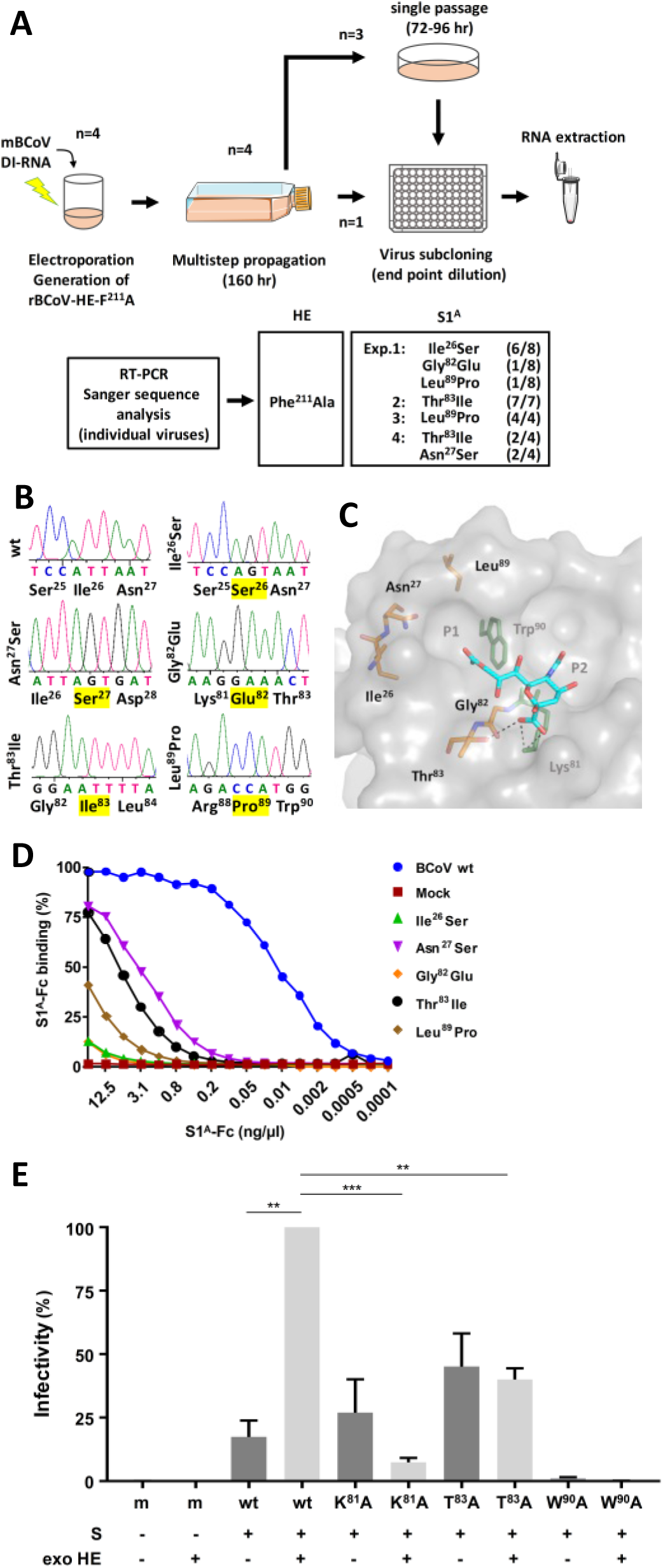
Site-directed mutagenesis of BCoV HE by targeted recombination; loss of HE lectin activity selects for second-site mutations in domain 1^A^ of the spike protein. (*A*) Schematic outline of four independent experiments, depicting each step from targeted RNA recombination and virus rescue to virus purification and genetic analysis of the resultant clonal populations by RT-PCR and Sanger sequencing. The number of virus clones, found to contain a particular S1^A^ mutation, relative to the total number of virus clones analyzed are given (between parenthesis). (*B*) Relevant portions of Sanger DNA sequencing chromatograms of the S1^A^ coding region in recombinant wildtype BCoV and in cloned rBCoV-HE-Phe^211^Ala derivatives. Amino acid substitutions marked in yellow. (*C*) Second-site mutations in S1^A^ locate in close proximity of the RBS. Close-up of the BCoV S1^A^ RBS (in surface representation; PDB:4H14), with 9-*O*-Ac-Sia (in sticks, colored by element; oxygen, red; nitrogen, blue; carbons, cyan) modeled in the RBS^32^, showing the locations of the mutations. Key elements of the RBS (hydrophobic pockets P1 and P2, and the side chains of RBS residues Lys^81^ and Trp^90^ in sticks, colored green) are indicated. Side chains and connecting main chains of RBS mutations also shown in sticks, but with carbon colored orange. Predicted hydrogen bonds between the Sia carboxylate moiety and the side chains of Lys^81^ and Thr^83^ shown as black dashed lines. (*D*) The S1^A^ mutations strongly reduce binding to 9-*O*-Ac-Sia. Mutant S1^A^-Fc fusion proteins in twofold serial dilutions, starting at 2.5 µg/well, were tested by sp-LBA for their binding to BSM relative to that of wildtype S1^A^-Fc. Binding expressed in percentages with maximum binding of wildtype S1^A^-Fc set to 100%. (*E*) Infectivity of VSV particles, pseudotyped with low affinity BCoV S variants, is inhibited rather than promoted by soluble ‘exogenous’ sialate-*O*-acetylesterase. HRT18 cells were inoculated with equal amounts of G-deficient VSV particles, pseudotyped with BCoV S or mutants thereof, either with (+) or without (-) soluble exogenous sialate-*O*-acetylesterase (‘exo HE’) added to the inoculum. ‘Infectivity’ expressed in RLUs in cell lysates at 18 h p.i., normalized to those measured for VSV-S^wt^. The data shown are averages from three independent experiments, each of which performed with technical triplicates. SDs and significant differences, calculated by Welch’s unequal variances *t* test, are indicated (\*\**P* ≤ 0.01; \*\*\**P* ≤ 0.001).

Generating BCoV-Rluc derivatives defective in HE lectin function proved more cumbersome. To abolish the HE RBS, we substituted Phe^211^, which is key to ligand binding^17^(**sFig. 1B**), by Ala via two nucleotide substitutions to block reversion. Mutant viruses were recovered eventually, but, in 3 out of 4 successful trials, a multistep 160-hr rescue did not suffice and an additional 72-96 hr blind passage was required (**Fig. 1A**).

Sequence analysis of clonal virus populations, obtained by endpoint dilution, confirmed the presence of the HE Phe^211^Ala substitution in all cases. Surprisingly, the purified viruses all suffered single site mutations in S, clustering in domain S1^A^ (aa 15-302) in proximity of the RBS (**Figs. 1A-C; sFig. 2**). Two of the trials yielded multiple S variants and some variants -Thr^83^Ile and Leu^89^ Pro-emerged independently in separate experiments (**Fig. 1A**). The mutations map to three distinct S RBS elements (**Fig. 1C, sFig. 2C**; nomenclature according to^32^). Ile^26^Ser and Asn^27^Ser locate in the S1^A^ β1 element within the N-terminal L1-β1-L2 segment (aa 15-33) that walls pocket P1; P1 is crucial for ligand binding as it accommodates the all-important sialate-9-*O*-acetyl moiety^32,36^. Moreover, in the OC43 S cryo-EM holostructure, the Asn^27^ side chain hydrogen bonds with the 9-*O*-acetyl carbonyl^36^. Leu^89^Pro in S1^A^ element 3_10_1 is immediately adjacent to Trp^90^. The latter is arguably the most critical residue in the RBS as its indole side chain separates the P1 and P2 pockets, and its replacement precludes receptor-binding and virus infectivity^32^. Finally, Gly^82^Glu and Thr^83^Ile substitutions occurred in S1^A^ element β5 that interacts with the sialate carboxylate through hydrogen bonding with Lys^81^ and Thr^83^.

**Fig. 2.**
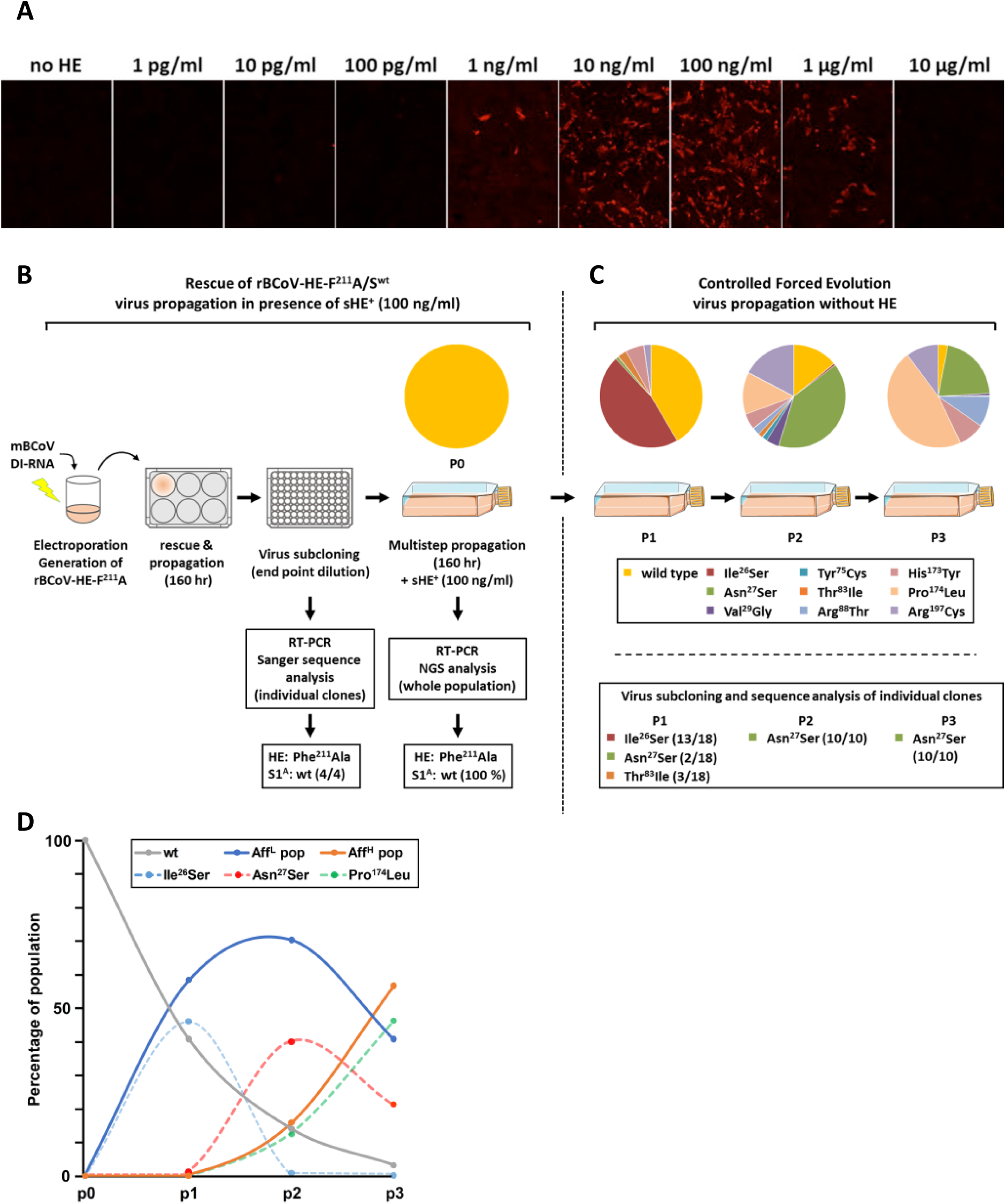
Stable propagation and controlled directed evolution of rBCoV-HE-F^211^ A. (*A*) rBCoV-HE-F^211^A propagation and spread is enhanced by soluble exogenous sialate-*O*-acetylesterase. mBCoV-infected LR7 cells, donor RNA-transfected to generate rBCoV-HE-F^211^A, were seeded on HRT18 cell monolayers to rescue recombinant viruses with cell culture supernatants supplemented with purified BCoV HE^+^-Fc at concentrations indicated. Cell supernatants, harvested 120 hr after seeding, were inoculated onto HRT18 cells grown on glass coverslips. After a single infectious cycle (12 hr p.i.), infected cells were identified by immunofluorescence assay. Infected cells stained red. (*B*) Stable maintenance of wildtype S protein in rBCoV-HE-F^211^A in the presence of exogenous HE and (*C*) forced evolution in the absence thereof. Visual representation of experimental procedures and findings. Generation of recombinant rBCoV-HE-F^211^A by targeted recombination was as in Fig. 1, but with rescue, cloning and virus amplification steps performed with tissue culture media supplemented with 100ng/ml exogenous HE-Fc. Rescued virus was purified by endpoint dilution. Individual clonal populations were characterized for HE and S1^A^ master sequences by extracting viral genomic RNA from the cell culture supernatant followed by RT-PCR and Sanger sequencing. One clonal population was used to grow a p0 stock of rBCoV-HE-F^211^A and HE and S1^A^ diversity was assessed by next gen illumina sequence analysis (NGS). The virus was used to inoculate 5 × 10^6^ HRT18 cells at an MOI of 0.005 TCID50/cell and serially passaged. Cell culture supernatants were harvested at 120 hr p.i. and virus diversity was assessed by subcloning and genetic analysis of purified viruses as in Fig. 1. In addition, viral RNA was extracted, and diversity determined by RT-PCR amplification and NGS. Frequencies of S1^A^ variants are presented in pie charts with individual color coding as indicated. (*D*) Propagation of rBCoV-HE-F^211^A quasispecies selects for variants with (near) wildtype S affinity. Stylized graph representation depicting the emergence and decline of viral variants during serial passage of rBCoV-HE-F^211^A in the absence of exogenous HE. Changes in the frequencies of variants with wildtype S and groups of variants with low affinity S (sum of Ile^26^Ser, Asn^27^Ser, Val^29^Gly, Thr^83^Ile, His^173^Tyr and Arg^197^Cys) and high affinity S (sum of Arg^88^Thr and Pro^174^Leu) are depicted with solid lines, colored in gray, blue and orange, respectively. Those of individual S variants, Ile^26^Ser, Asn^27^Ser and Pro^174^Leu, are shown in dashed lines and colored light blue, red and green, respectively.

As measured by solid phase lectin binding assay (sp-LBA) with S1^A^-Fc fusion proteins and bovine submaxillary mucin (BSM) as ligand, all mutations significantly reduced S binding to 9-*O*-Ac-Sia albeit to widely different extents. S1^A^-Fc binding affinities of the mutants were 500-fold (Asn^27^Ser) to more than 10.000-fold (Ile^26^Ser; Gly^82^Glu) lower than that of parental BCoV S1^A^-Fc (**Fig. 1D**).

Loss of the HE lectin RBS would both abolish HE-assisted virion attachment to 9-*O*-Ac-Sia receptor determinants as well as reduce virion-associated receptor-destroying sialate-*O*-acetylesterase activity towards clustered glycotopes^11^. The findings suggest that the HE-Phe^211^Ala substitution creates a fitness defect that can be alleviated not by increasing but by dramatically reducing the affinity of the S RBS. Thus, the defect apparently is caused by loss of virion-associated RDE activity rather than by loss of overall virion avidity.

### Loss of HE lectin function exerts a fitness cost by affecting reversible virion attachment

In principle, downregulation of HE esterase activity consequential to loss of lectin function, could affect virus propagation at two stages of the infectious cycle, namely virus release, which would require depletion of intracellular and cell surface receptor pools, and (pre)attachment. We recently reported that G-deficient vesicular stomatitis (VSV) virions pseudotyped with wildtype BCoV S require exogenous HE for efficient infection^32^. S1^A^ mutations that reduce S affinity inhibit infection, but, as we now show, only when HE is present. VSV virions pseudotyped with low affinity mutant S proteins were less reliant on or even inhibited by exogenous HE with decreasing S RBS affinity (**Fig. 1E**). The findings provide direct proof that the S1^A^ mutations act at the level of virion attachment and support the notion that the S1^A^ mutations, selected for in HE-defective rBCoVs, restore reversibility of receptor binding by lowering S affinity and thereby protect against inadvertent attachment to decoy receptors.

### HE lectin-deficient recombinant BCoVs are genetically stable when grown in the presence of exogenous receptor-destroying enzyme

To test whether loss of virion-associated RDE activity in rBCoV-HE-F^211^A/S^wt^/Rluc might be compensated for by adding exogenous soluble HE to the culture medium, we seeded infected/transfected LR7 cells onto HRT18 cell monolayers, supplemented the cell culture supernatant with BCoV HE-Fc^17^ to final concentrations of 1 pg to 10 μg/ml, and allowed infection to proceed for 120 hr. While in the absence of HE-Fc there was no sign of virus propagation as detectable by IFA, concentrations of exogenous sialate-*O*-acetylesterase as low as 1 ng/ml to up to 1 μg/ml promoted virus growth (**Fig. 2A**).

To determine whether these conditions would allow isolation of rBCoV-HE-F^211^A without mutations in S1^A^, we performed targeted recombination and rescued recombinant viruses by 160 hr multistep propagation as before, but now with culture supernatant supplemented with 100 ng/ml HE-Fc (**Fig. 2B**). Sanger sequence analysis of RT-PCR amplicons showed that all viruses cloned by endpoint dilution of the 160-hr stock (n=4) coded for mutant HE-Phe^211^Ala in combination with wildtype S1^A^. To assess the stability of clonal rBCoV-HE-F^211^A/S^wt^/Rluc, the virus population resulting from a subsequent 120-hr amplification in the presence of exogenous HE-Fc was analyzed by Next-Generation Sequencing (NGS), which allows for the detection of low frequency mutants. Sequence variation in HE and S1^A^ was distributed randomly and did not exceed background levels (<0.15%). More than 99.5% of the viruses coded for HE-Phe^211^Ala, while preserving parental type S1^A^ (**Fig. 2B**).

### Loss of HE lectin function gives rise to mixed virus population with competition and cooperativity among S affinity variants

With a clonal, virtually pure stock of rBCoV-HE-F^211^A/S^wt^ available, we performed controlled forced evolution experiments. The virus was serially passaged involving three consecutive 120-hr multistep propagation rounds in HRT18 cells but now in the absence of exogenous HE-Fc, with the initial infection at multiplicity of infection (MOI) of 0.005 (**Fig. 2C**). In trial 1, viral titers in passage 1 (p1) increased only slowly to 3 × 10^4^ and 2 ×10^4^ TCID50/ml (measured with or without exogenous HE-Fc, respectively). The withdrawal of exogenous RDE during viral passage immediately selected for mutations in S1^A^. Virus cloning by endpoint dilution of the 120-hr p1 sample yielded S RBS mutants Asn^27^Ser, Thr^83^Ile and Ile^26^Ser (**Fig. 2C**) – all three of which had been seen before (**Fig. 1A**). NGS analysis revealed the true complexity of the p1 population (**Fig. 2C**) and identified two additional S1^A^ variants with substitutions -His^173^Tyr and Arg^197^Cys-more distal from the RBS (**sFigs. 2D, 3A, B**). His^173^Tyr also reduced the relative binding affinity of S1^A^-Fc albeit less dramatically than the other mutations, namely by 30-fold (**Table 1**). The Arg^197^Cys mutation seemingly falls in a separate category as it reduced S1^A^-Fc expression levels by more than 90% suggestive of defective folding (**sFig. 3C**). Apparently, aberrant disulfide-bonding causes most of the fusion protein to be retained in the ER with only a minor, presumably properly folded fraction slipping through to become secreted.

**Table 1.** Virus composition in passages p0 through p3 of controlled forced evolution experiment 1. ^1^Percental occurrence of BCoV S1^A^ mutations and ^2^their relative binding affinities as measured by equilibrium endpoint solid phase binding assay with S1^A^-Fc fusion proteins with that of parental BCoV S1^A^-Fc (‘wildtype’) set at 1.0. S1^A^ variants Thr^22^Ile and Asn^27^Tyr emerged only in experiment 2 (see sFig. 5), but their affinities relative to that of wildtype S1^A^ are shown for comparison. (NA, not applicable).

|  | Frequency in population<sup>1</sup> |  |  |  |  |
| --- | --- | --- | --- | --- | --- |
| S1<sup>A</sup> | p0 | p1 | p2 | p3 | rAff<sup>2</sup> |
| wildtype | 100 | 40.90 | 13.90 | 3.00 | 1.0 |
| Ile<sup>26</sup>Ser | 0 | 45.83 | 0.70 | 0.03 | 0.0000625 |
| Asn<sup>27</sup>Ser | 0 | 0.94 | 39.80 | 21.09 | 0.004 |
| Val<sup>29</sup>Gly | 0 | 0 | 4.10 | 0.65 | 0.008 |
| Tyr<sup>75</sup>Cys | 0 | 0 | 1.60 | 0.02 | ND |
| Thr<sup>83</sup>Ile | 0 | 2.90 | 1.60 | 0.21 | 0.002 |
| Arg<sup>88</sup>Thr | 0 | 0 | 2.30 | 9.69 | 0.25 |
| His<sup>173</sup>Tyr | 0 | 5.77 | 5.00 | 8.10 | 0.03 |
| Pro<sup>174</sup>Leu | 0 | 0 | 13.00 | 46.72 | 0.5 |
| Arg<sup>197</sup>Cys | 0 | 2.24 | 17.25 | 10,.27 | 0.25 |
| Thr<sup>22</sup>Ile | NA | NA | NA | NA | 0.015 |
| Asn<sup>27</sup>Tyr | NA | NA | NA | NA | 0.002 |

**Fig. 3.**
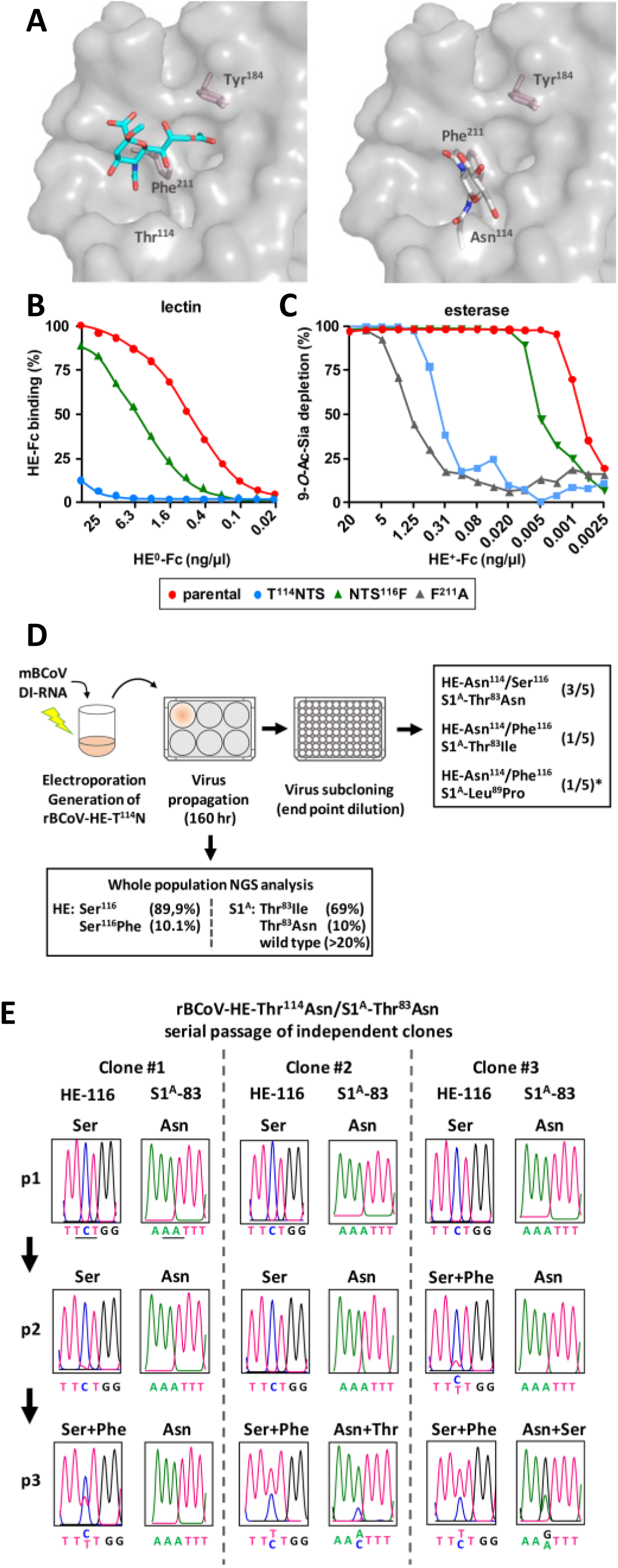
BCoV S and HE co-evolve to restore functional balance and optimal virion avidity. (*A*) Introduction of an N-glycosylation site at the rim of the HE RBS. Side-by-side close ups of the BCoV HE holostructure^17^ (PDB 3CL5) in surface representation with 9-*O*-Ac-Sia (in sticks, colored by element; oxygen, red; nitrogen, blue; carbons, cyan) bound to the RBS (left) or without the ligand and with the Thr^114^Asn substitution and N-linked glycan modelled in the RBS (in sticks, colored by element as above but with carbons in white) (right). Modelling performed by superpositioning of the OC43 strain NL/A/2005 HE structure (PDB 5N11) and confirmed in Coot. HE RBS key residues are indicated, with side chains of Tyr^184^ and Phe^211^ shown in sticks. (*B*) Loss of HE lectin function upon introduction of an N-glycosylation site through HE-Thr^114^Asn substitution and partial restoration through a second-site Ser^116^Phe mutation. Sp-LBA with serial dilutions of enzyme-inactive HE^0^-Fc as in Fig. 1D. (*C*) Consequences for HE esterase activity towards clustered ligands. On-the-plate receptor depletion assay with bovine submaxillary mucin as substrate as in^11^. The assay was performed with 2-fold serial dilutions of enzymatically active HE^+^-Fc and residual 9-*O*-Ac-Sia measured by sp-LBA with a fixed amount of HE^0^-Fc. (*D*) Visual representation of experimental procedures and findings as in Figs. 1 and 2. Note that of five virus clones purified by endpoint dilution, the identity of one isolate could not be established and was deduced from subsequent propagation experiments as explained in the text (marked with *; see also **Fig. 4**). (*E*) Serial passage of rBCoV-HE-Thr^114^Asn/S1^A^-Thr^83^Asn selects for successive mutations in HE and S to restore their function to near wildtype levels. Results are shown for three independent isolates. Viral RNA extracted from tissue culture supernatants collected at the end of each passage was characterized by RT-PCR and Sanger DNA sequencing. Relevant portions of Sanger DNA sequencing chromatograms are presented to show changes in the master sequence of the virus populations at the coding sequence for the N-glycosylation site (HE codon 116) and for the site of the low affinity S mutation selected initially (S1^A^ codon 83).

All in all, the p1 population was comprised for virtually 100% of HE-Phe^211^Ala mutants, 40% of which in combination with parental BCoV S, the remaining 60% with second-site mutations in S1^A^ (**Fig. 2C**; **Table 1**). Of the latter, the ultra-low affinity variant S1^A^-Ile^26^Ser was the most abundant at 46% and the Asn^27^Ser variant the least at less than 1%. However, upon a subsequent round of 120-hr multistep propagation, the tables were turned with S1^A^-Asn^27^Ser now comprising almost 40% of the p2 population and the Ile^26^Ser variant reduced to 0.7%. In addition, four other S1^A^ variants emerged. One of these had a mutation in S1^A^ RBS loop L1, Val^29^Gly, and a relative binding affinity close to that of the Asn^27^Ser mutant (**Fig. 2C**; **Table 1**). We also identified at position 75 a second S1^A^ Cys-substitution mutant, which like Arg^197^Cys, presumably disrupts the RBS through aberrant disulfide bonding (**sFig. 3**). Remarkably, two other S1^A^ variants arose with mutations -Arg^88^Thr and Pro^174^Leu-that affected the relative binding affinity only modestly to 0.25 and 0.5 of that of wildtype S1^A^-Fc, respectively (**Table 1**). Even more remarkably, upon further passage these mutants increased to dominate the p3 population, effectively outcompeting variants with low affinity spikes as well as those with parental spikes (**Fig. 2C, D**). However, when the p1, p2 and p3 stocks were cloned by endpoint dilution in the absence of exogenous HE-Fc, only virus variants with low affinity mutations in S1^A^ were isolated (**Fig. 2C; Table 1**). Strikingly, from the p3 stock, the S1^A^-Asn^27^Ser variant was isolated exclusively against all odds (10/10 tested; p < 10^−6^) when calculated purely from its frequency in the population (21%). Conversely, virus cloning by endpoint dilution in the presence of exogenous HE-Fc yielded high affinity S1^A^ mutants Pro^174^Leu (5/11) and Arg^88^Thr (4/11), parental virus rBCoV-HE-Phe^211^Ala/S^wt^ (1/11), and intermediate S affinity variant His^173^Tyr (1/11).

Notably, the conditions selected not only for mutations in S but also in HE. Variants with an Ala^211^Val substitution in HE emerged in p2, rising to 17% of the p2 end population, to stabilize around this frequency in p3. As a result of this mutation, HE lectin affinity was regained albeit to levels solely detectable by high-sensitivity nanobead hemagglutination assay, while esterase activity towards clustered glycotopes in BSM increased 4-fold as compared to HE-Phe^211^Ala, but still remained 125-fold lower than that of wildtype HE (**sFig. 4**). Apparently, the increase in HE function, minor as it may be, provides a selective advantage, but apparently one that benefits both low and high affinity S variants, because the mutation was found in cloned viruses of either type.

**Fig. 4.**
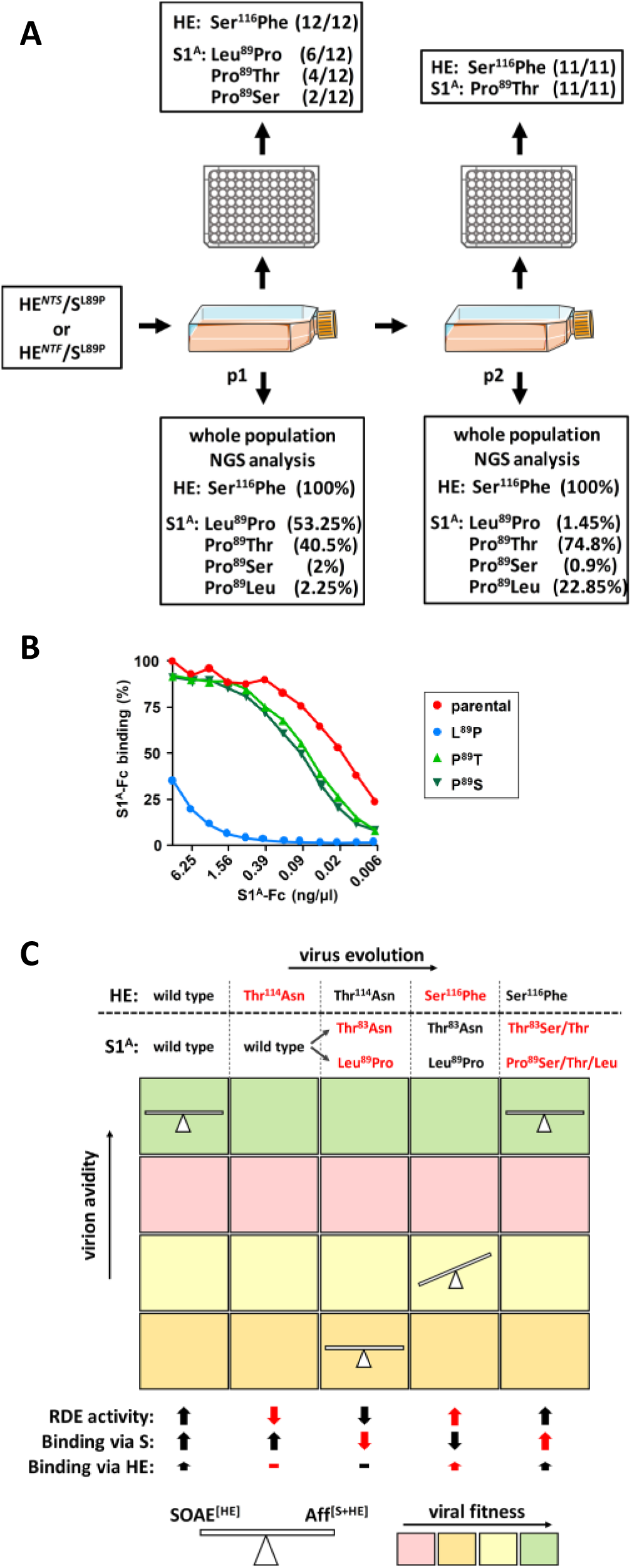
Serial passage of rBCoV-HE-T^114^A selects for successive mutations in HE and S to optimize HE/S functional balance and overall virion avidity. (*A*) Continued serial passage of rBCoV-HE-T^114^N+S^116^F/S1^A^-L^89^P. Schematic outline of the experiment and presentation of results of NGS analysis and genetic characterization of subcloned variants as in Figs. 1, 2 and 3. (*B*) Re-substitution of S1^A^-Leu^89^Pro by Thr or Ser restores S1^A^ binding to near wildtype levels. Sp-LBA as in Fig. 1D. (*C*) BCoV is under selective pressure for an optimal functional balance between virion attachment and catalysis-driven release as well as for optimal virion avidity. Schematic summary of the evidence for HE-S coevolution and our interpretation thereof in a two-dimensional chart. The course of evolution of rBCoV-HE-Thr^114^Asn (direction indicated by arrow) is shown for two types of low affinity S escape variants (Thr^114^Asn and Leu^89^Pro) with the succession of mutations in HE and S selected for during serial passage (top) brought in relation to (i) overall virion avidity, as mediated by S and HE, on the Y-axis from low to high as indicated by the arrow, (ii) viral fitness, color-coded from low (pink) to high (green) as indicated in the color legend at the bottom, (iii) the effect of the mutations on the function of S (attachment) and HE (receptor-destroying enzyme (RDE) activity and attachment) as indicated by thick arrows (arrows pointing up, near wildtype activity; arrows pointing down, decreased function; ?, total loss of function; the size difference between arrows for S and HE reflect the difference in their contribution to virion binding; to indicate the effect of newly emerging mutations corresponding arrows for function are colored red) and (iv) their effect on HE-S functional balance (as indicated by the position of the scale).

### Loss of HE lectin function selects for virus swarms with low affinity S escape mutants promoting the emergence and propagation of high-affinity S variants

To corroborate our observations, the controlled forced evolution experiment was repeated (**sFig. 5**). As compared to the first trial, there was a much faster population built-up already in p1 at 120 hr p.i. with final titers reaching 4 × 10^8^ and 3.4 × 10^7^ TCID50/ml, when measured in the presence or absence of exogenous HE-Fc, respectively. Surprisingly, in stark contrast to trial 1, the trial 2 p1 population was comprised for about 94% of viruses expressing wildtype BCoV S. Less than 6% consisted of variants with mutations in S1^A^, four of low receptor binding affinity (Thr^22^Ile, Asn^27^Tyr, Val^29^Gly, His^173^Tyr), one of near-wildtype binding affinity (Pro^174^Leu), and, with the exception of Thr^22^Ile, all at positions seen before (**Tables 1, 2**). Consistent with our previous findings, however, virus purification through endpoint dilution in the absence of exogenous HE-Fc yielded low affinity mutants (11/11 tested) exclusively (**sFig. 5**). If the variants in the trial 2 p1 population were all of equal replicative fitness under the conditions applied, the odds of this result would be less than 1.10^−12^.

**Table 2.** Summary of S1^A^ mutations identified in rBCoVs upon i. introduction of a HE-Phe^211^Ala substitution and virus rescue by straight forward targeted recombination (E1 through E4, see also Fig. 1), ii. passage of rBCoV-HE-F^211^A/S^wt^ in the absence of exogenous HE (E5 and E6) or iii. introduction of a HE-Asn^114^Thr substitution and subsequent viral passage (E7).

| **S1<sup>A</sup> residue** | **E1** | **E2** | **E3** | **E4** | **E5** | **E6** | **E7** |
| --- | --- | --- | --- | --- | --- | --- | --- |
| **Thr-22** |  |  |  |  |  | **Ile** |  |
| **Ile-26** | **Ser** |  |  |  | **Ser** |  |  |
| **Asn-27** |  |  |  | **Ser** | **Ser** | **Tyr** |  |
| **Val-29** |  |  |  |  | **Gly** | **Gly** |  |
| **Tyr-75** |  |  |  |  | **Cys** |  |  |
| **Gly-82** | **Glu** |  |  |  |  |  |  |
| **Thr-83** |  | **Ile** |  | **Ile** | **Ile** |  | **Ile/Asn** |
| **Arg-88** |  |  |  |  | **Thr** |  |  |
| **Leu-89** | **Pro** |  | **Pro** |  |  |  | **Pro** |
| **His-173** |  |  |  |  | **Tyr** | **Tyr** |  |
| **Pro-174** |  |  |  |  | **Leu** | **Leu** |  |
| **Arg-197** |  |  |  |  | **Cys** |  |  |

At first glance, the two trials would seem to differ in their outcomes. We offer, however, that the results are in fact consonant and that the main difference is in the speed with which the virus populations increased and evolved. There is an inherent stochastic element to the experimental approach and whether the developing quasispecies undergoes slow track (experiment 1) or fast track evolution (experiment 2) is likely dependent on the time of advent of the first mutant virus and its properties, for instance whether it is an ultralow (like Ile^26^Ser) or low affinity variant (like Asn^27^Ser). The findings allow for several conclusions. (i) They confirm and firmly establish that loss of HE lectin function selects for mutations in S1^A^ that reduce S receptor-binding affinity and virion avidity. (ii) The possibilities to reduce the affinity of the S RBS through single site mutations are finite. In several independent experiments, substitutions in S1^A^ occurred at a limited number of positions albeit not necessarily by the same residue. For example, Asn^27^ was replaced both by Ser and Tyr. (iii) The mutations that reduce S affinity fall into different categories. Most map within or in close proximity of the RBS to affect receptor-ligand interaction directly. Others, like His^173^Tyr and Pro^174^Leu, are more distal from the RBS and apparently affect ligand binding indirectly through long range conformational effects. A third type of mutations, quasi-random Cys substitutions, apparently disrupt S1^A^ folding by promoting non-native disulfide-bonding. While an Arg^197^Cys substitution strongly decreased secretion of the S1^A^-Fc fusion protein, the biosynthesis and intracellular transport of native trimeric spikes was seemingly affected to lesser extent. At least, the uptake of S-Arg^197^Cys into VSV pseudotypes was not noticeably impaired as compared to that of wildtype S (**sFig. 3E**). Still, the mutation did alter the infectivity of the pseudotyped particles making them less dependent on exogenous HE-Fc (**sFig. 3F**) presumably by reducing the avidity of S trimers through local S1^A^ misfolding and consequential disruption of the RBS in one or more monomers. (iv) Perhaps most surprisingly, quasispecies developed in which loss of HE lectin function was compensated at the level of the viral population with minority low affinity variants, constituting less than 6% of the swarm, not only sustaining the replication of high affinity variants but actually allowing the latter to flourish and amplify to become the majority phenotype.

### S and HE proteins co-evolve to attain functional balance and optimal virion avidity

Among the first mutations fixed upon zoonotic introduction and early emergence of OC43, was a HE-Thr^114^Asn substitution, which created a glycosylation site at the rim of the lectin domain RBS^11^ (**Fig. 3A**). Glycans attached to HE Asn^114^ hamper binding to 9-*O*-Ac-Sia through steric hindrance, causing a 500-fold reduction in HE avidity (**Fig. 3B**) and a 125-fold in sialate-*O*-acetylesterase-activity, respectively (**Fig. 3C**). We introduced the HE Thr^114^Asn substitution in BCoV, expecting that the glycosylation site would be rapidly lost through any of several single-nucleotide restorative mutations in HE. Indeed, NGS analysis of the virus swarm arising after targeted recombination showed the glycosylation site to be destroyed but only in 10% of the population and exclusively by Ser^116^Phe substitution (**Fig. 3D**). This mutation partially restores HE receptor binding and receptor destruction to 0.125 and 0.17 of that of wildtype HE, respectively (**Fig. 3B, C**). In the vast majority of viruses, the newly introduced HE glycosylation site was retained and, instead, S1^A^ mutations that reduced S affinity were selected again, with S-RBS Thr^83^ replaced either by Ile (69%) -as seen before (**Figs. 1A, 2C; Table 1**)- or by Asn (10%) (**Fig. 3D**). The latter mutation reduces S1^A^ affinity to 0.008 of that of wildtype.

Virus cloning by endpoint dilution yielded, in three out of five isolates, S1^A^-Thr^83^Asn variants with the newly introduced N-glycosylation site in HE intact (HE-Thr^114^Asn). Furthermore, a single S1^A^-Thr^83^Ile variant was isolated, but this virus in addition had the N-glycosylation site in HE destroyed (HE-Thr^114^Asn/Ser^116^Phe) (**Fig. 3D**). The observations led us to entertain the possibility that the mutations in S1^A^ and HE did not occur independently and that, even in viruses expressing low affinity spikes, partially restorative mutations in HE would yet provide a selective advantage. To test this, the clonal S1^A^-Thr^83^Asn/HE-Thr^114^Asn variants were serially passaged. All three viruses independently lost HE Asn^114^ glycosylation over time and, saliently, through Ser^116^Phe substitution exclusively. Even more remarkably, with HE-Ser^116^Phe mutants gaining dominance, variants emerged that had restored S affinity to (near)wildtype through substitution of S1^A^-Asn^83^ either by Thr or by Ser (**Fig. 3E**).

For one of the five clonal populations obtained by endpoint dilution, we unfortunately failed to determine its genotype for technical reasons. From the NGS analysis of the p1 population, we deduced that the starting mutant must have been a low affinity S1^A^-Leu^89^Pro variant that, like the S1^A^- Thr^83^Asn/Ile variants described above, quickly lost the HE-Asn^114^ glycan through an HE-Ser^116^Phe substitution. Oddly enough, the Leu^89^Pro substitution had not been detected by NGS in the pre-cloning virus stock. Note, however, that this mutation had been selected before twice independently in trials with rBCoV-HE-F^211^A (**Table 2**). Possibly, it arose spontaneously during end point dilution procedure. Be that as it may, its *in vitro* evolution proved informative (**Fig. 4A**). NGS analysis of a passage p1 population, resulting from 120-hr multistep propagation, showed that 100% of the viruses coded for HE-Thr^114^Asn/Ser^116^Phe in combination with S1^A^-Pro^89^ (53.3%), - Thr^89^ (40.5%), or -Ser^89^ (2%). Note that the relationship between these variants and the course of evolution –from Leu^89^ in the parental recombinant virus to Pro and from Pro to Thr or Ser– is evident from the codon sequences (CTA→C***C***A→***T/A***CA) and that the Thr^89^ and Ser^89^ substitutions restored S RBS affinity almost to that of wildtype RBS (**Fig. 4B**). All three variants - S1^A^-Pro^89^, -Thr^89^ and -Ser^89^, were readily cloned and isolated by standard endpoint dilution, and propagated independently without a requirement for exogenous RDE. p1 also contained a minor population of viruses with parental S1^A^, presumably regenerated from S1^A^-Pro^89^, which as for the S1^A^-Thr^89^ and Ser^89^ variants would have required only a single nucleotide substitution (CCA→C***T***A). Apparently, with HE lectin function partially restored, viruses that regained (near) wildtype S affinity had a selective advantage. At the end of passage p2, S1^A^-Pro^89^ variants had dwindled to less than 1.5%, S1^A^ -Thr^89^ had become dominant at 75% and viruses with parental S1^A^-Leu^89^ had rapidly risen from 2.25% in p1 to 23% (**Fig. 4A**).

In summary, the introduction of a glycosylation site in the HE lectin domain that reduced receptor-binding affinity and, thereby, reduced sialate-*O*-acetylesterase activity towards clustered glycotopes, triggered a series of successive mutations in S and HE. Thus, the data directly demonstrate HE-S co-evolution. Moreover, the findings suggest that virions are under selective pressure not only to balance receptor-binding and receptor-destroying activities in apparent relation to cell-surface receptor-densities, but also, within these constrains, to maximize virion avidity.

### Cell culture adapted BCoV and OC43 strains differ in their set point of the S/HE balance

The impact of loss of function mutations in the BCoV HE lectin domain was unexpected, because this defect seems well tolerated by the prototype OC43 laboratory strain USA/1967. In HRT18 cells, it grows to titers comparable to those of BCoV reference strain Mebus. Next gen sequencing of OC43 stocks revealed heterogeneity, but no indications for the existence of low S affinity minority variants that would support replication of majority high S affinity viruses. Also, clonal virus populations obtained by end point dilution (10/10) all conformed to the S1^A^ master sequence. We offer that instead OC43-USA/1967 may have reached a viable S/HE balance compatible with efficient *in vitro* propagation through adaptations in S that reduced receptor-binding affinity and/or altered receptor fine-specificity. When measured by solid phase assay with bivalent S1^A^-Fc fusion proteins, binding of the S protein of OC43 USA/1967 to bovine submaxillary mucin, containing both mono- and di-*O*-acetylated α2,6-sialoglycans, is 16 to 32-fold lower than that of BCoV-Mebus^32^. sp-LBA with BSM preparations, selectively depleted for either 9-*O-* or 7,9-di-*O*-Sias, showed that BCoV S, like BCoV HE^28^, preferentially binds to 7,9-di-*O*-Ac-Sias (**sFig. 6A**). OC43 USA/1967 S1^A^ may not share this preference. Apparently due to its low affinity, detectable binding to BSM was lost upon depletion of either type of Sia (**sFig. 6A**). Moreover, even though BCoV-Mebus S preferably binds to 7,9-di-*O*-Ac-Sia, monovalent one-on-one binding of the BCoV S1^A^ domain to α2,6-linked 9-*O*-acetylated Sia is still 3-fold stronger than that of OC43-USA/1967 as measured by biolayer interferometry (**sFig. 6B**). On a cautionary note, the isolation and complex passage history of OC43-USA/1967^43,44^ entailed several passages in human tracheal organ culture, suckling mouse brain and many rounds of replication in cultured cells^9^, which would have given the virus ample opportunity to adapt to the *in vitro* conditions. Thus, the binding characteristics of its spike may not faithfully reflect those in circulating field variants. Indeed, OC43 variants in sputum samples, contrary to the OC43 USA/1967, replicate in airway epithelial cell cultures but not in tissue culture cells^45^.

## DISCUSSION

### Co-evolution and functional interdependence of embecovirus S and HE proteins

Our findings demonstrate that in the prototypic β1CoV BCoV the envelope proteins S and HE are functionally entwined and co-evolve. We posit that the same holds for other members of the species *Betacoronavirus-1*, including its zoonotic descendant human coronavirus OC43 and related viruses of swine, rabbits, dogs and horses, as well as for other *Embecovirus* species, most prominently among which human coronavirus HKU1. The data lead us to conclude that the respective activities of S and HE in receptor-binding and catalysis-driven virion elution are balanced to ensure dynamic reversible virion attachment and, thereby, efficient virus propagation. In consequence, for the viruses listed above, the roles of S and HE during natural infection cannot be understood in isolation but must be considered in unison.

Using a reverse genetics-based forced evolution approach with BCoV as a model system, we showed that loss of HE lectin function causes an offset in S-HE balance, practically incompatible with virus propagation and spread. With the HE lectin domain as modulator of esterase activity, mutations that decrease or abolish HE RBS affinity reduce virion-associated sialate-*O*-acetylesterase activity towards clustered glycotopes on hypervalent glycoconjugates^11^ such as are present in the mucus and glycocalyx in natural tissues and on the surface of cultured cells. The extent of the resultant defect is such that compensatory second-site mutations in S are selected for that dramatically reduce S RBS affinity, apparently to restore reversibility of binding as an escape ticket from inadvertent virion attachment to non-productive sites.

The single-amino acid mutations in receptor-binding domain S1^A^ were limited to a finite number of positions, either within or proximal to the RBS to directly affect protein-ligand interactions, or more distal to reduce RBS affinity through long range effects or by disrupting local folding through aberrant disulfide-bonding. Whereas the parental recombinant viruses, defective in HE lectin function but with wildtype S RBS affinity, require an external source of receptor-destroying enzyme for propagation, their progeny escape mutants regained propagation-independence by lowering S affinity.

In expanding clonal populations of HE-defective rBCoV-HE-F^211^A, propagated in the absence of exogenous receptor-destroying enzyme activity, viruses with reduced S affinity gained a selective advantage initially. Upon prolonged passage, however, quasispecies developed in which loss of HE lectin function was compensated at the population level. Variants that combined the HE-Phe^211^Ala mutation with (near) wildtype affinity S proteins increased to dominate the swarm at least numerically. Still, these high affinity S variants for their proliferation were strictly reliant on minority low affinity S variants. This relationship extends beyond cooperativity and group selection described for other systems^46–52^ and amounts to a state of dependency. We propose that the virions of the low affinity minority variants provide aid by serving as a source of exogenous sialate-*O*-acetylesterase activity. They themselves evade decoy receptors through enhanced reversibility of virion attachment, but this phenomenon increases their motility -whether by sliding diffusion or binding-rebinding-causing them to deplete cell surface 9-*O*-Ac-Sias, decoy receptors and functional receptors alike. With increasing concentrations of low affinity virions in the culture supernatant, high affinity variants would profit progressively, whereas falling cell surface receptor densities would put the low affinity viruses increasingly at a disadvantage.

The forced evolution trials performed with rBCoV-HE-F^211^A were restricted in course and outcome by design, because full reversion would require simultaneous mutation of two adjacent nucleotides. Moreover, the crucial role of the Phe^211^ in ligand binding (**sFig. 1**) obviates conservative substitutions ^17^. Although rBCoV-HE^A211V^ variants did emerge in two separate experiments, this mutation only marginally increases HE RBS affinity and sialate-*O*-acetylesterase activity.

In contrast to the HE-Phe^211^Ala mutation, the deleterious effect of N-glycosylation at HE-Asn^114^ can be reversed, completely or partially, through various single-nucleotide substitutions in codons 114 and 116 and would therefore more readily allow for compensatory mutations also in HE. Indeed, serial passage of the rBCoV-HE-Thr^114^Asn resulted in a succession of mutations alternatingly in HE and S. The order of appearance of these mutations and their effect on protein function indicated that they were not fixed to merely restore the balance between attachment and catalysis-driven virion elution. The HE-Thr^114^Asn substitution initially selected for second-site mutations that reduced S affinity (Thr^83^Ile, Thr^83^Asn and Leu^89^Pro), but with propagation thus recovered, derivatives rapidly emerged with increased HE lectin and esterase activity through a Ser^116^Phe mutation. Apparently, this created an HE-S disbalance that in turn favored the selection of viruses with revertant mutations in S that raised S RBS affinity again to wildtype (Thr^83^ → Ile _→_ Thr; Leu^89^ → Pro _→_ Leu) or near wildtype levels (Thr^83^ → Ile → Ser; Leu^89^ → Pro_→_ Thr/Ser). Conjointly, our findings indicate that through an initial sharp reduction in overall avidity, compensatory to loss of HE function, virus particles regained the capacity of eluding non-productive attachment to decoy receptors, but at a fitness penalty. The decrease in S RBS affinity would predictably lower the specific infectivity of virus particles through a decrease in productive host cell attachment. The rapid selection of the HE-Ser^116^Phe mutation in a low-affinity S background can thus be understood to have increased virion avidity, albeit through HE and rather than through S. HE does have a dual function after all and in influenza viruses C and D as well as in murine coronavirus-1, it is a receptor-binding protein first and foremost^22,35,53–56^. Of note, the partial Ser^116^Phe reversion of HE consistently seen in multiple independent experiments, suggests that a return to (near) wildtype lectin and esterase activity along with a low affinity S would have tipped the scale too much towards catalytic virion release. We posit that in addition to an optimal balance between receptor-binding and receptor-destruction, the system strives towards optimal virion avidity (**Fig. 4C**).

Under natural circumstances, the set-point of the S/HE balance would be tailored to conditions met in the target tissues of the intact host. The spontaneous loss of HE lectin function in OC43 and HKU1 may thus be understood to have arisen through convergent evolution as an adaptation to the sialoglycan composition of the mucus in the human upper respiratory tract and that of the glycocalyx of the respiratory epithelia^11^. This change, which would predictably reduce virion-associated receptor-destruction and hence decrease virion elution/increase or prolong virion attachment, might have been selected for by low density occurrence of 9-*O*-Ac-sialoglycans in the human upper airways. In accordance, limited tissue array analyses with HE-based virolectins suggested that these sugars are not particularly prevalent in the human respiratory tract and by far not as ubiquitous as in the gut^28^. However, full understanding of how the S-HE balance was reset in OC43 and HKU1 upon their zoonotic introduction and why awaits further analysis of the binding properties and ligand fine-specificity of the S proteins of naturally occurring variants as well as more quantitative and comprehensive inter-host comparative analyses of airway sialoglycomes. As an added complication, virion particles encounter widely different circumstances while traversing the mucus layer, at the epithelial cell surface, during local cell-to-cell dissemination, and during transmission. It is an open question whether this selects for majority phenotypes that can cope individually and independently with each of these different conditions by striking an uneasy compromise with regard to HE/S balance and overall virion avidity, or whether there is loco-temporal selection for swarms of variants that collectively allow the virus population as a whole to overcome each hurdle.

### Similarities between embeco- and influenza A viruses point to common principles of virion-sialoglycan receptor-usage

The embecovirus HE gene originated from a horizontal gene transfer event, presumably with an influenza C/D-like virus as donor^17,57^. Like the orthomyxovirus hemagglutinin-esterase fusion proteins, the newly acquired coronavirus HE protein provided the acceptor virus with an opportunity to reversibly bind to 9-*O*-Ac-sialoglycans^26^. This in turn would seem to have prompted a shift in the receptor-specificity of S through adaptations in S1^A^ that created a 9-*O*-Ac-Sia binding site *de novo* so that virions could now attach to these receptor determinants also via S. The embecoviruses thus adopted a strategy of receptor usage entailing a concerted and carefully fine-tuned activity of two envelope proteins that is unique among coronaviruses, but uncannily similar to that of influenza A viruses. In the latter, the hemagglutinin (HA), as a pendant of S, mediates binding to either α2,3- or α2,6-linked sialosides, while the neuraminidase, like HE, is a receptor-destroying enzyme with a substrate fine-specificity that closely matches HA ligand preference^58^. For influenza A virus, the existence and biological relevance of a functional balance between receptor-binding and receptor destruction is well recognized^59–62^. This balance is critical for receptor-associated virus motility through the mucus and at the cell surface^63–68^. Complete or partial loss of NA activity -whether invoked spontaneously, through reverse genetics or by viral propagation in the presence of NA inhibitors-selects for mutations around the HA receptor-binding pocket that reduce HA affinity^60,69,70^. Furthermore, as proposed here for HE, NA contributes to virion attachment and even compensates for loss of virion avidity in mutant viruses with reduced HA affinity^71^. Different from HE, NA may do so via its catalytic pocket which doubles as a Sia-binding site^72^. However, NA also possesses a second Sia binding site^73,74^, which like the HE lectin domain, regulates NA activity and which, in further analogy, is conserved or lost in apparent correlation with host tropism^75–77^. Finally, among many other similarities to embecoviruses, influenza A variants with different set points in their HA-NA functional balance may cooperate to support their propagation in cultured cells^48^. Our observations establish that there are common principles of virion-sialoglycan interactions that prompted convergent evolution of β1CoVs and influenza A viruses. Although these two groups of viruses essentially differ in genome type and replication strategy, envelope proteins, and receptors, they seem to be subject to the same rules of engagement with respect to dynamic receptor-binding, the differences between them constituting variations on a theme. This implies that observations made for the one system are informative for the other. Perhaps more importantly, insight into the overriding principles of virus-glycan interactions may open avenues to common strategies for antiviral intervention.

## Methods

### Cells and viruses

Human rectal tumor (HRT) 18 (ATCC® CCL-244™) and mouse LR7^41^ cells were maintained in Dulbecco’s modified Eagle’s medium (DMEM) containing 10% fetal calf serum (FCS), penicillin (100 IU/ml) and streptomycin (100 µg/ml). BCoV strain Mebus and OC43 strain USA/1967, purchased from the American Type Culture Collection (ATCC), were propagated in HRT18 cells.

### Reverse genetics through targeted recombination

A reverse genetics system based on targeted RNA recombination was developed for BCoV strain Mebus essentially as described^15,41,78^. Using conventional cloning methods, RT-PCR amplicons of the 5’-terminal 601 nts and 3’-terminal 9292 nts of the BCoV strain Mebus genome (reference Genbank sequence U00735.2) were fused and cloned in plasmid pUC57, downstream of a T7 RNA polymerase promotor and upstream of a 25-nt poly(A) tract and a *Pac*I site, yielding pD-BCoV1. From this construct, BCoV ORF 4a was deleted (nts 27740-27853) and replaced by the *Renilla* luciferase (Rluc) gen, yielding pD-BCoV-Rluc. A second pD-BCoV1 derivative, pD-mBCoVΔHE, was created by replacing the coding sequence for the ectodomain of BCoV S (nts 23641-27433) by the corresponding MHV-A59 sequence and by deleting the BCoV HE gene (nts 22406-23623). The nucleotide sequences of pD-BCoV1, pD-BCoV-Rluc and pD-mBCoVΔHE, determined by bidirectional Sanger sequence analysis, were deposited in Genbank (accession codes: XXX).

To generate a recombinant chimeric acceptor virus, mBCoVΔHE, HRT18 cells were infected with BCoV-Mebus at a MOI of 10 TCID50/cell and trypsinized and re-suspended in PBS. An aliquot of this suspension, containing 1.5 × 10^6^ cells in 0.8 ml, was mixed with capped synthetic RNA that had been produced by *in vitro* transcription using the mMESSAGE mMACHINE™ T7 Transcription Kit (Thermo fisher) with *Pac*I-linearized mBCoVΔHE vector as template. The mixture was subjected to two consecutive electrical pulses of 850 V at 20 μF with a Gene Pulser II electroporator (Bio-Rad) and the cells were then seeded on a confluent monolayer of LR7 feeder cells in a 35mm dish. Incubation was continued at 37°C, 5% CO_2_ for 18 hours post transfection until wide-spread cytopathic effect (CPE) was apparent. The cell culture supernatant was harvested and cleared by low speed centrifugation at 1200 rpm, and mBCoVΔHE was purified by end-point dilution and used to generate stocks for future usage in LR7 cells.

To generate luciferase-expressing rBCoVs with the BCoV HE and S genes reconstituted, i.e. rBCoV^wt^ or rBCoV-HE-Phe^211^Ala, LR7 cells, infected with mBCoVΔHE at MOI 5, were electroporated as described above with synthetic RNA transcribed from pD-BCoV-Rluc and derivatives thereof. The infected and transfected cells were then seeded on HRT18 cell monolayers in 35-mm plates for up to 160 hr. For rescue and propagation of rBCoV-HE-Phe^211^Ala without second-site mutations in S, the cell culture supernatants were supplemented with 100 ng/ml of BCoV HE-Fc protein^17^. After 5-7 days of incubation at 37°C, samples of the cell culture supernatants were tested for infectivity by transferring them to HRT18 cell monolayers grown on 12-mm glass coverslips in 15.6-mm wells. Incubation was continued for 12 hr after which the cells were fixed with paraformaldehyde and immunofluorescence staining was performed with polyclonal antiserum from a BCoV-infected cow.

### Virus titration, purification and characterization of viral populations

mBCoV was titrated and cloned by endpoint dilution on LR7 cells with cytopathic effect as read-out. rBCoVs were titrated and cloned in HRT18 cells. To identify infected wells, cell supernatants were analyzed by hemagglutination assay with rat erythrocytes^26^ and by *Renilla* luciferase assay (Dual-Luciferase® Reporter Assay System, Promega). Titers were calculated by the Spearman-Kaerber formula. Clonal virus populations were characterized by isolating viral RNA from 150 µl aliquots of the cell culture supernatant with the NucleoSpin® RNA Virus kit (MACHEREY-NAGEL) followed by conventional RT-PCR and bidirectional Sanger sequence analysis.

### Controlled forced evolution experiments

Confluent HRT18 monolayers (5 × 10^6^ cells) grown in 25-cm^2^ flasks, were inoculated with rBCoV-HE-F^211^A at an MOI 0.005 in PBS for 1 hr at 37°C. The cells were washed three times with PBS to remove residual exogeneous HE-Fc and incubation was continued in DMEM + 10% FCS at 37°C, 5% CO_2_ for 120 hr post infection (pi) with samples collected every 24 hours (passage 1). Subsequent 120-hr passages were performed by adding 10 µL of supernatant to new cultures of HRT18 cells in 25-cm^2^ flasks.

### Expression and purification of HE-Fc and S1^A^-Fc proteins

BCoV HE, either enzymatically-active (HE^+^) or rendered inactive through a Ser^40^Ala substitution (HE^0^), and OC43 S1^A^ were expressed as Fc fusion proteins in HEK293T cells and purified from the cell supernatant by protein A affinity chromatography as detailed^17,32^. Monomeric S1^A^ was obtained by on-the bead thrombin cleavage^32^. pCD5-BCoVHE-T-Fc vectors^17^ encoding mutant BCoV HE derivatives were constructed with the Q5® Site-Directed Mutagenesis Kit per the instructions of the manufacturer.

### Pseudovirus entry assays

The production of BCoV S-pseudotyped VSV-ΔG particles, their characterization by Western blot analysis, and infectivity assays in HRT18 cells were as described^32^.

### Solid-phase lectin binding assay (sp-LBA)

sp-LBA was performed as described^32^ with bovine submaxillary mucin (BSM; Sigma-Aldrich), coated to 96-Well Maxisorp® microtitre ELISA plates (Nunc, 0.1 µg BSM per well), serving as a ligand. Binding assays were performed with 2-fold serial dilutions of HE^0^-Fc, S1^A^-Fc, or mutated derivatives thereof. Receptor-destroying esterase activities of soluble HEs were measured by on-the-plate 9-*O*-Ac-Sia depletion assays as described^11,28^.

### Hemagglutination assay (HAA)

HAA was performed with rat erythrocytes (*Rattus norvegicus* strain Wistar; 50% suspension in PBS). Standard HAA was done with two-fold serial dilutions of HE^0^-Fc proteins (starting at 25 ng/well) as described^17^. High sensitivity nanoparticle HAA (NP-HAA) was performed as in^32,79^. Briefly, self-assembling 60-meric nanoparticles, comprised of lumazine synthase (LS), N-terminally extended with the immunoglobulin Fc-binding domain of the *S. aureus* protein A, were complexed with HE^0^-Fc proteins at a 1:0.6 molar ratio for 30 min on ice. The HE^0^-Fc-loaded nanoparticles were then 2-fold serially diluted and mixed 1:1 (vol/vol) with rat erythrocytes (0.5% in PBS). Incubation was for 2 hr at 4°C after which HAA titers were read.

### NGS analysis

Viral RNA from culture supernatants was isolated as described above. HE and S1^A^ coding regions from viral genome of different virus populations were obtained by RT-PCR with primer sets HE_F_ 5’-TTAGATTATGGTCTAAGCATCATG-3’ and HE_R_ 5’-TTAGATTATGGTCTAAGCATCATG-3’, S1^A^_F_ 5’-ACCATGTTTTTGATACTTTTA-3’ and S1^A^_R_ 5’-AGATTGTGTTTTACACTTAATCTC-3’, respectively. Amplicons were processed in the NGSgo® workflow for Illumina according to the Instructions for Use (Edition 4), except that the fragmentation was prolonged to 40 min at 25°C (protocol 3A). Briefly, amplicons were subjected to fragmentation and adapter ligation using NGSgo-LibrX (GenDx). Size selection and clean-up of the samples was performed with SPRI beads (Machery-Nagel). Unique barcodes were ligated to each sample using NGSgo-IndX (GenDx), after which all samples were pooled and subsequently purified with SPRI beads, resulting in a library of fragments between ∼400 and 1000 bp. The DNA fragments were denatured and paired-end sequenced on a MiSeq platform (Illumina) using a 300 cycle kit (V2). FASTQ files were analyzed in NGSengine® (GenDx), which aligned the reads to the reference sequences of HE and S (reference Genbank sequence U00735.2 for BCoV strain Mebus, and NC_006213.1 for OC43 strain USA/1967). For the characterization of each virus sample, amplicons from five independent RT-PCR reactions were analyzed in parallel and mutation frequencies were determined by averaging the results from these five replicates.

## Author contributions

Y.L., W.L. and R.J.d.G. conceived the study and designed research; Y.L., W.L., D.K. and A.C.S.v.B. performed research; E.R. and H.M.v.S. performed NGS analysis; Y.L., W.L., Z.L., D.K., A.C.S.v.B., E.R., G.J.P.H.B., F.J.M.v.K., B.J.B., E.G.H., H.M.v.S. and R.J.d.G. analyzed data; W.L., Z.L. and G.J.P.H.B. provided and synthesized reagents; Y.L. and R.J.d.G. wrote the paper. All authors discussed the results and W.L., Z.L., G.J.P.H.B., F.J.M.v.K., B.J.B., E.G.H. and H.M.v.S. commented on the manuscript.

## Declaration of Interests

The authors declare no competing interests.

