## supplementary information for "Coronavirus hemagglutinin-esterase and spike proteins co-evolve for functional balance and optimal virion avidity"

### Slide 1
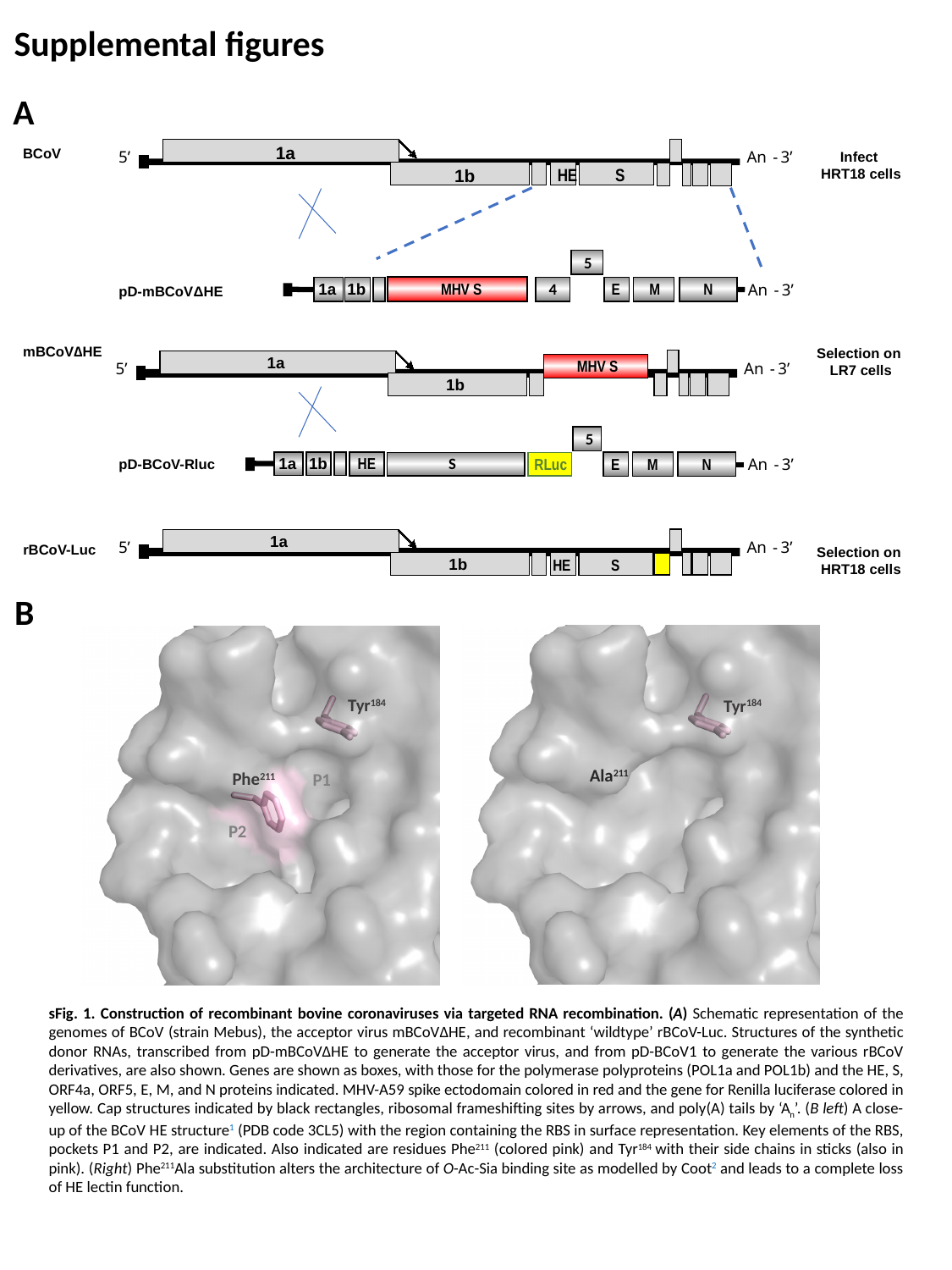

Supplemental figures
A
1a
5’
An
-
3’
HE
S
1b
An
-
3’
An
-
3’
BCoV
Infect
HRT18 cells
pD-mBCoVΔHE
mBCoV∆HE
Selection on
LR7 cells
MHV S
5
1a
1b
HE
S
RLuc
E
N
M
pD-BCoV-Rluc
1a
5’
An
-
3’
1b
HE
S
rBCoV-Luc
Selection on
HRT18 cells
5
1a
1b
MHV S
4
E
M
N
1a
5’
An
-
3’
1b
B
Tyr^184^
Tyr^184^
Ala^211^
Phe^211^
P1
P2
sFig. 1. Construction of recombinant bovine coronaviruses via targeted RNA recombination. (A) Schematic representation of the genomes of BCoV (strain Mebus), the acceptor virus mBCoV∆HE, and recombinant ‘wildtype’ rBCoV-Luc. Structures of the synthetic donor RNAs, transcribed from pD-mBCoVΔHE to generate the acceptor virus, and from pD-BCoV1 to generate the various rBCoV derivatives, are also shown. Genes are shown as boxes, with those for the polymerase polyproteins (POL1a and POL1b) and the HE, S, ORF4a, ORF5, E, M, and N proteins indicated. MHV-A59 spike ectodomain colored in red and the gene for Renilla luciferase colored in yellow. Cap structures indicated by black rectangles, ribosomal frameshifting sites by arrows, and poly(A) tails by ‘A_n_’. (B left) A close-up of the BCoV HE structure^1^ (PDB code 3CL5) with the region containing the RBS in surface representation. Key elements of the RBS, pockets P1 and P2, are indicated. Also indicated are residues Phe^211^ (colored pink) and Tyr^184^ with their side chains in sticks (also in pink). (Right) Phe^211^Ala substitution alters the architecture of O-Ac-Sia binding site as modelled by Coot^2^ and leads to a complete loss of HE lectin function.

### Slide 2
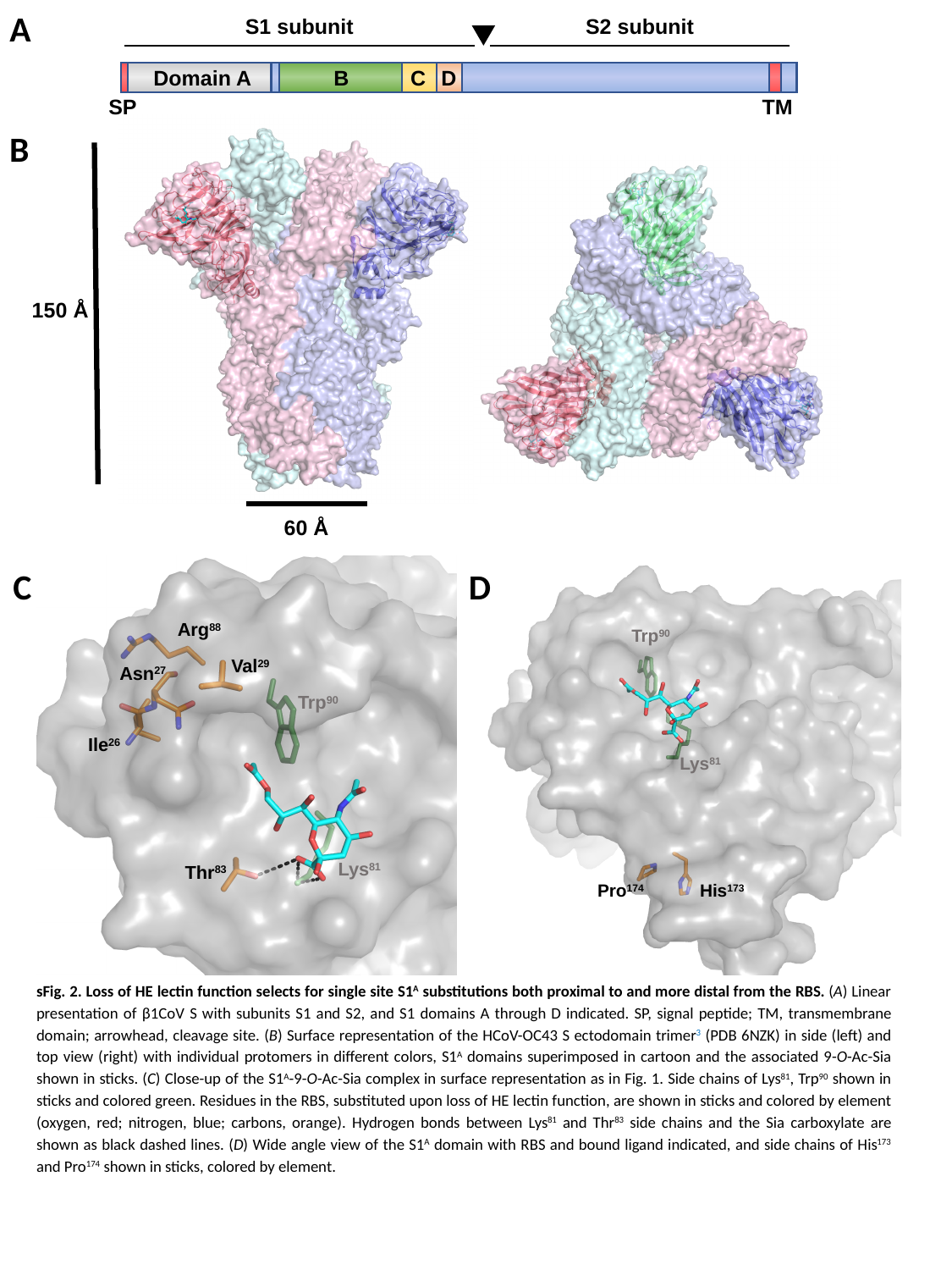

A
S1 subunit
S2 subunit
Domain A
B
C
D
SP
TM
B
150 Å
60 Å
Trp^90^
Lys^81^
Pro^174^
His^173^
Arg^88^
Val^29^
Asn^27^
Trp^90^
Ile^26^
Lys^81^
Thr^83^
C
D
sFig. 2. Loss of HE lectin function selects for single site S1^A^ substitutions both proximal to and more distal from the RBS. (A) Linear presentation of β1CoV S with subunits S1 and S2, and S1 domains A through D indicated. SP, signal peptide; TM, transmembrane domain; arrowhead, cleavage site. (B) Surface representation of the HCoV-OC43 S ectodomain trimer^3^ (PDB 6NZK) in side (left) and top view (right) with individual protomers in different colors, S1^A^ domains superimposed in cartoon and the associated 9-O-Ac-Sia shown in sticks. (C) Close-up of the S1^A^-9-O-Ac-Sia complex in surface representation as in Fig. 1. Side chains of Lys^81^, Trp^90^ shown in sticks and colored green. Residues in the RBS, substituted upon loss of HE lectin function, are shown in sticks and colored by element (oxygen, red; nitrogen, blue; carbons, orange). Hydrogen bonds between Lys^81^ and Thr^83^ side chains and the Sia carboxylate are shown as black dashed lines. (D) Wide angle view of the S1^A^ domain with RBS and bound ligand indicated, and side chains of His^173^ and Pro^174^ shown in sticks, colored by element.

### Slide 3
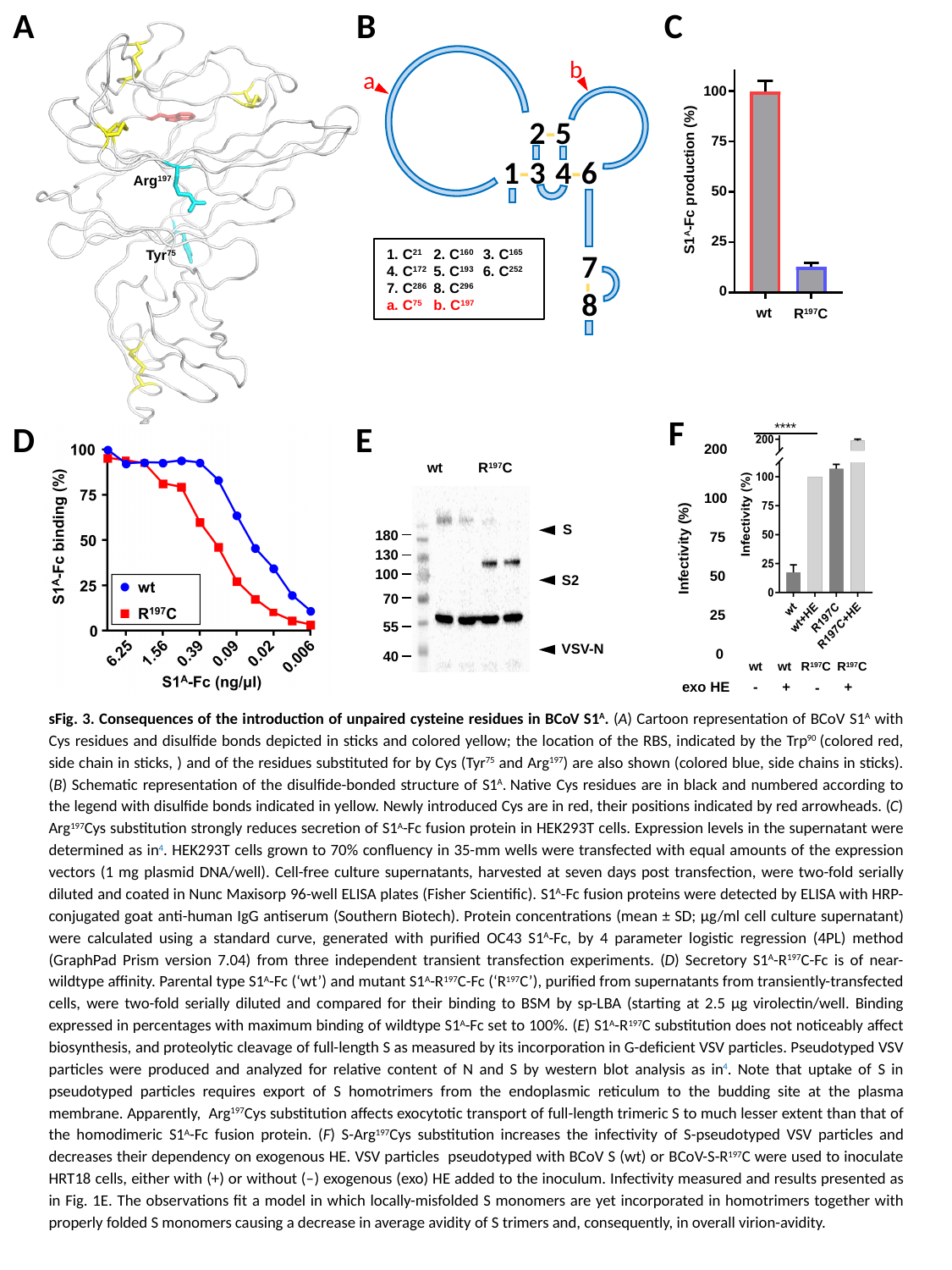

A
B
C
Arg^197^
Tyr^75^
100
75
50
25
0
S1^A^-Fc production (%)
wt
R^197^C
b
a
2-5
1-3
4-6
7
8
-
1. C^21^ 2. C^160^ 3. C^165^
4. C^172^ 5. C^193^ 6. C^252^
7. C^286^ 8. C^296^
a. C^75^ b. C^197^
F
****
200
100
75
Infectivity (%)
50
25
0
wt
wt
R^197^C
R^197^C
exo HE
-
+
+
-
D
E
wt
R^197^C
S
180
130
100
S2
70
55
VSV-N
40
sFig. 3. Consequences of the introduction of unpaired cysteine residues in BCoV S1^A^. (A) Cartoon representation of BCoV S1^A^ with Cys residues and disulfide bonds depicted in sticks and colored yellow; the location of the RBS, indicated by the Trp^90^ (colored red, side chain in sticks, ) and of the residues substituted for by Cys (Tyr^75^ and Arg^197^) are also shown (colored blue, side chains in sticks). (B) Schematic representation of the disulfide-bonded structure of S1^A^. Native Cys residues are in black and numbered according to the legend with disulfide bonds indicated in yellow. Newly introduced Cys are in red, their positions indicated by red arrowheads. (C) Arg^197^Cys substitution strongly reduces secretion of S1^A^-Fc fusion protein in HEK293T cells. Expression levels in the supernatant were determined as in^4^. HEK293T cells grown to 70% confluency in 35-mm wells were transfected with equal amounts of the expression vectors (1 mg plasmid DNA/well). Cell-free culture supernatants, harvested at seven days post transfection, were two-fold serially diluted and coated in Nunc Maxisorp 96-well ELISA plates (Fisher Scientific). S1^A^-Fc fusion proteins were detected by ELISA with HRP-conjugated goat anti-human IgG antiserum (Southern Biotech). Protein concentrations (mean ± SD; μg/ml cell culture supernatant) were calculated using a standard curve, generated with purified OC43 S1^A^-Fc, by 4 parameter logistic regression (4PL) method (GraphPad Prism version 7.04) from three independent transient transfection experiments. (D) Secretory S1^A^-R^197^C-Fc is of near-wildtype affinity. Parental type S1^A^-Fc (‘wt’) and mutant S1^A^-R^197^C-Fc (‘R^197^C’), purified from supernatants from transiently-transfected cells, were two-fold serially diluted and compared for their binding to BSM by sp-LBA (starting at 2.5 µg virolectin/well. Binding expressed in percentages with maximum binding of wildtype S1^A^-Fc set to 100%. (E) S1^A^-R^197^C substitution does not noticeably affect biosynthesis, and proteolytic cleavage of full-length S as measured by its incorporation in G-deficient VSV particles. Pseudotyped VSV particles were produced and analyzed for relative content of N and S by western blot analysis as in^4^. Note that uptake of S in pseudotyped particles requires export of S homotrimers from the endoplasmic reticulum to the budding site at the plasma membrane. Apparently, Arg^197^Cys substitution affects exocytotic transport of full-length trimeric S to much lesser extent than that of the homodimeric S1^A^-Fc fusion protein. (F) S-Arg^197^Cys substitution increases the infectivity of S-pseudotyped VSV particles and decreases their dependency on exogenous HE. VSV particles pseudotyped with BCoV S (wt) or BCoV-S-R^197^C were used to inoculate HRT18 cells, either with (+) or without ( ̶ ) exogenous (exo) HE added to the inoculum. Infectivity measured and results presented as in Fig. 1E. The observations fit a model in which locally-misfolded S monomers are yet incorporated in homotrimers together with properly folded S monomers causing a decrease in average avidity of S trimers and, consequently, in overall virion-avidity.

### Slide 4
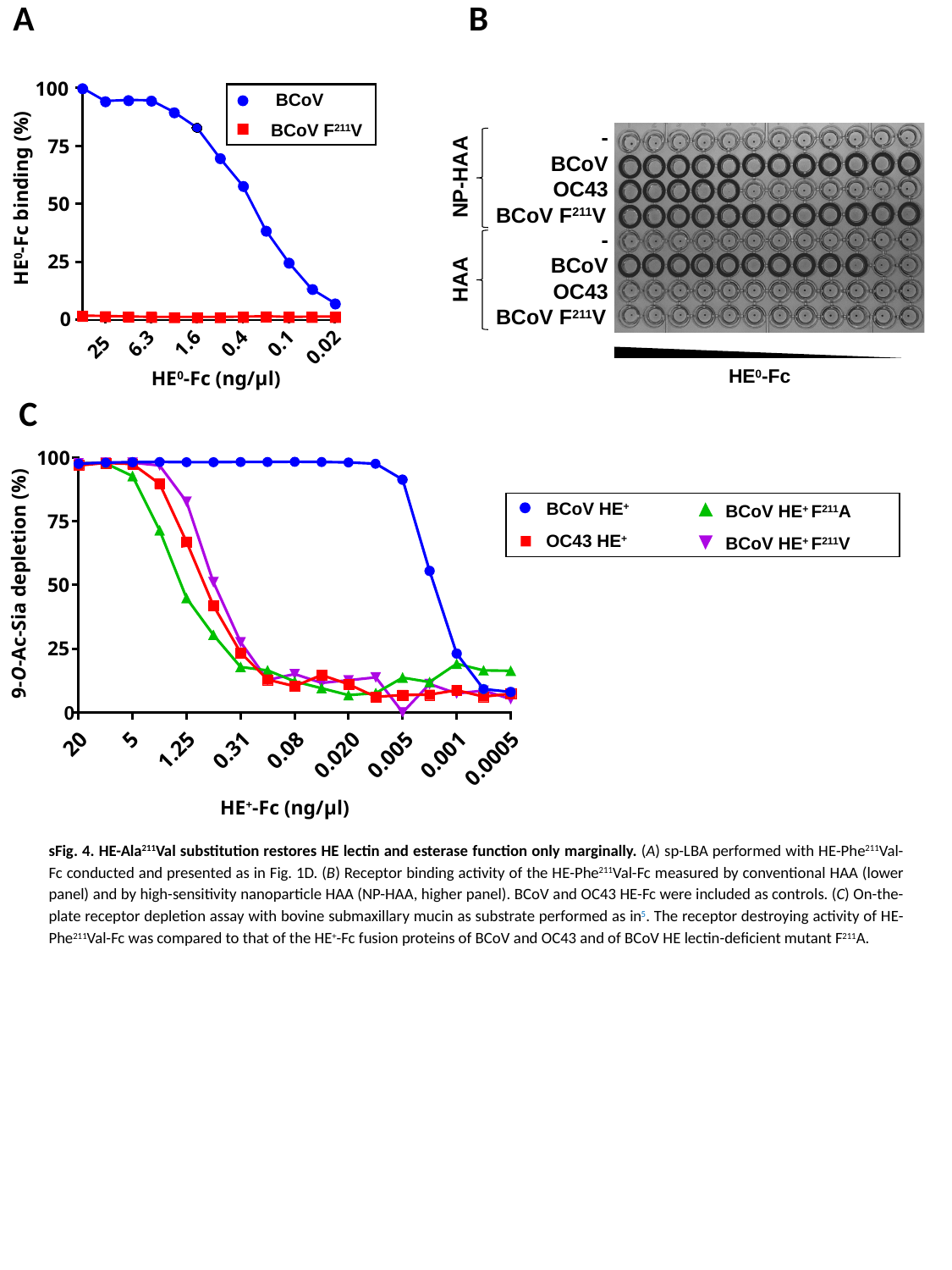

A
B
100
BCoV
BCoV F^211^V
75
50
25
HE^0^-Fc binding (%)
0
6.3
1.6
0.4
0.1
25
0.02
HE^0^-Fc (ng/μl)
-
BCoV
NP-HAA
OC43
BCoV F^211^V
-
BCoV
HAA
OC43
BCoV F^211^V
HE^0^-Fc
C
100
BCoV HE^+^ F^211^A
BCoV HE^+^
75
BCoV HE^+^ F^211^V
OC43 HE^+^
9-O-Ac-Sia depletion (%)
50
25
0
5
20
1.25
0.31
0.08
0.020
0.005
0.001
0.0005
HE^+^-Fc (ng/μl)
sFig. 4. HE-Ala^211^Val substitution restores HE lectin and esterase function only marginally. (A) sp-LBA performed with HE-Phe^211^Val-Fc conducted and presented as in Fig. 1D. (B) Receptor binding activity of the HE-Phe^211^Val-Fc measured by conventional HAA (lower panel) and by high-sensitivity nanoparticle HAA (NP-HAA, higher panel). BCoV and OC43 HE-Fc were included as controls. (C) On-the-plate receptor depletion assay with bovine submaxillary mucin as substrate performed as in^5^. The receptor destroying activity of HE-Phe^211^Val-Fc was compared to that of the HE^+^-Fc fusion proteins of BCoV and OC43 and of BCoV HE lectin-deficient mutant F^211^A.

### Slide 5
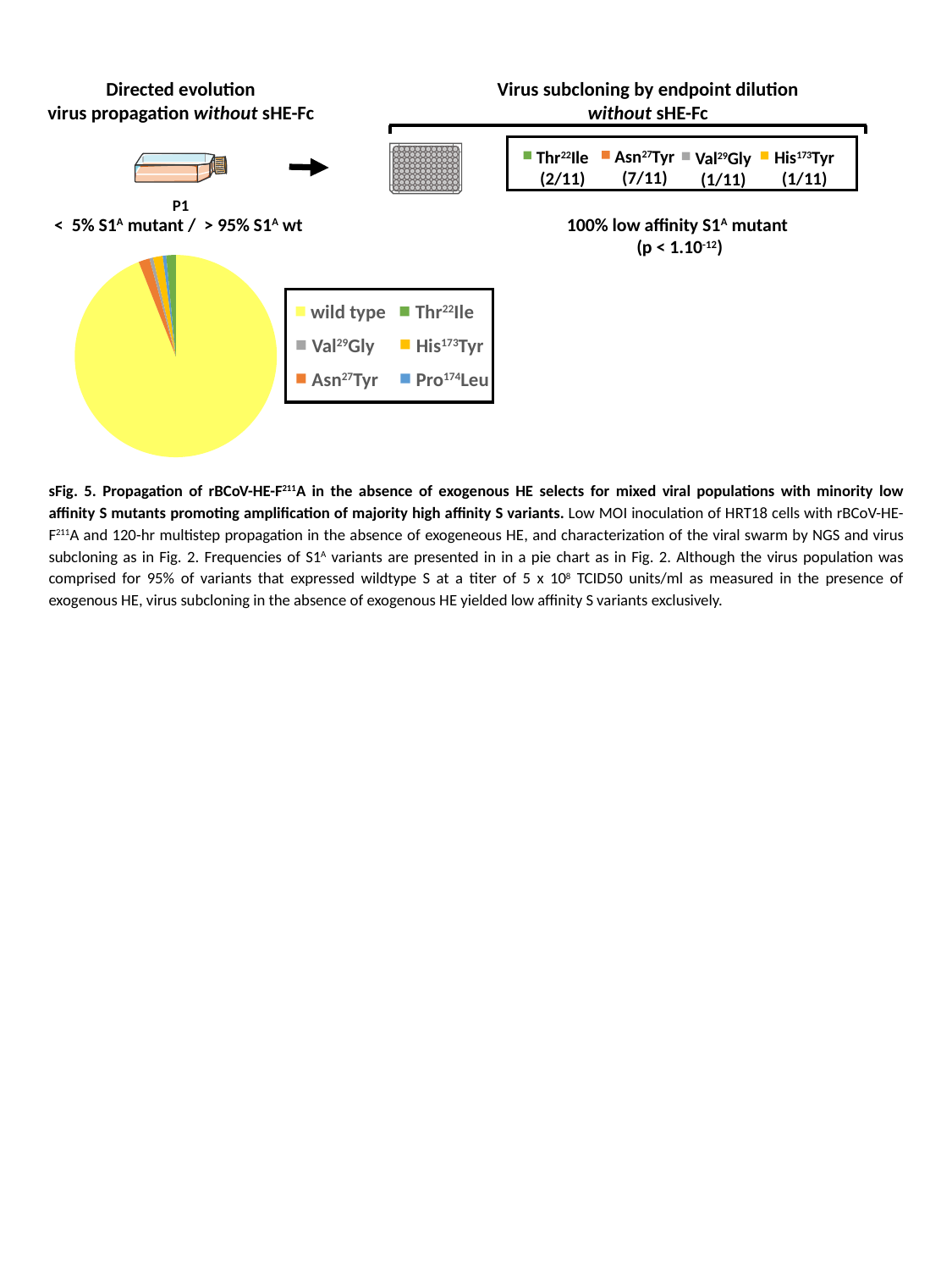

Directed evolution
virus propagation without sHE-Fc
Virus subcloning by endpoint dilution
without sHE-Fc
Asn^27^Tyr
(7/11)
Thr^22^Ile
(2/11)
His^173^Tyr
(1/11)
Val^29^Gly
(1/11)
P1
< 5% S1^A^ mutant / > 95% S1^A^ wt
100% low affinity S1^A^ mutant
(p < 1.10^-12^)
#### Chart
| Category | |
|---|---|
| wild type | 94.0 |
| Asn27 | 1.77 |
| Val29 | 0.58 |
| His173 | 1.55 |
| Pro174 | 0.55 |
| Thr22 | 1.52 |
Thr^22^Ile
wild type
His^173^Tyr
Val^29^Gly
Pro^174^Leu
Asn^27^Tyr
sFig. 5. Propagation of rBCoV-HE-F^211^A in the absence of exogenous HE selects for mixed viral populations with minority low affinity S mutants promoting amplification of majority high affinity S variants. Low MOI inoculation of HRT18 cells with rBCoV-HE-F^211^A and 120-hr multistep propagation in the absence of exogeneous HE, and characterization of the viral swarm by NGS and virus subcloning as in Fig. 2. Frequencies of S1^A^ variants are presented in in a pie chart as in Fig. 2. Although the virus population was comprised for 95% of variants that expressed wildtype S at a titer of 5 x 10^8^ TCID50 units/ml as measured in the presence of exogenous HE, virus subcloning in the absence of exogenous HE yielded low affinity S variants exclusively.

### Slide 6
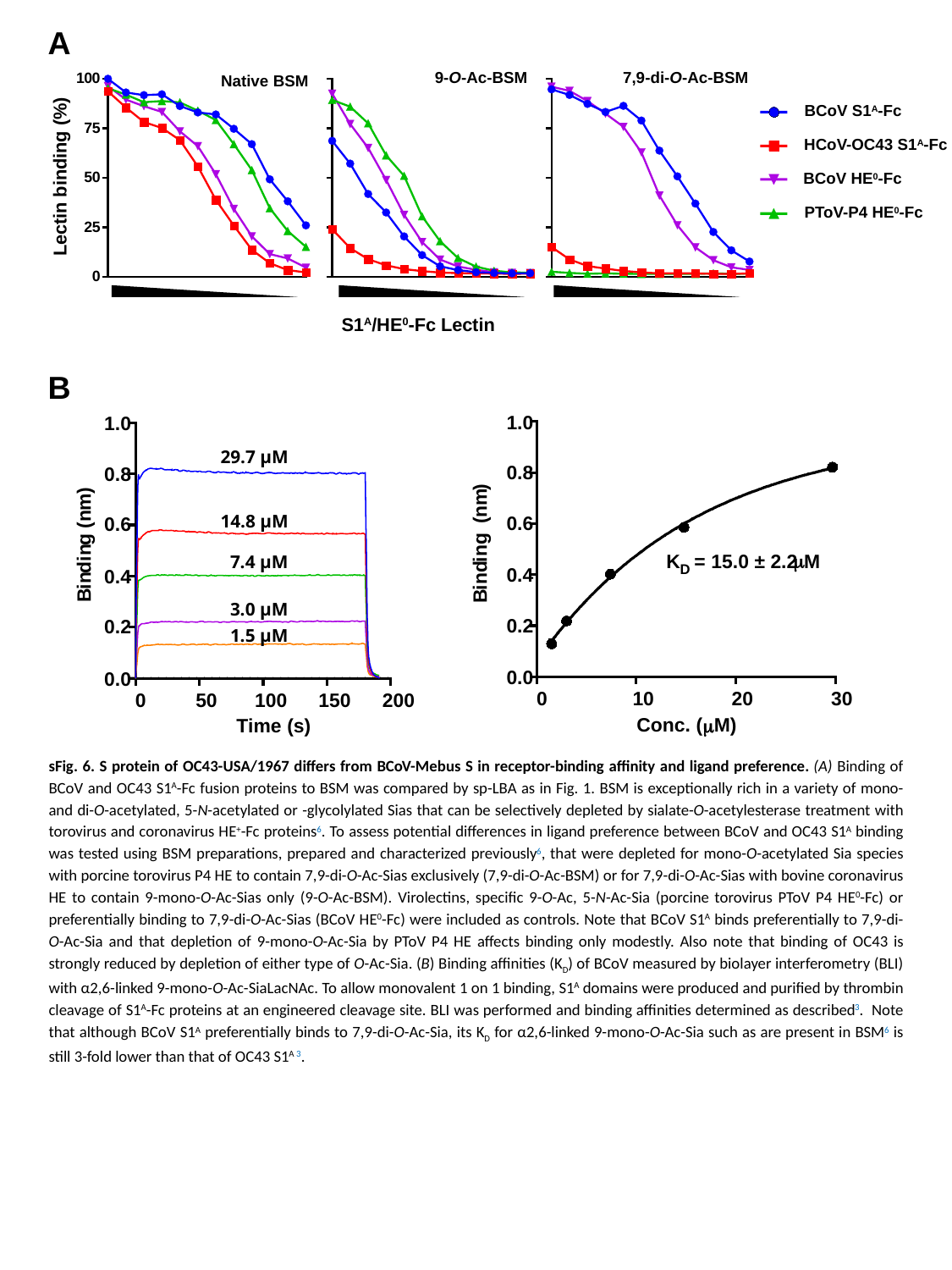

A
9-O-Ac-BSM
7,9-di-O-Ac-BSM
Native BSM
BCoV S1^A^-Fc
HCoV-OC43 S1^A^-Fc
Lectin binding (%)
BCoV HE^0^-Fc
PToV-P4 HE^0^-Fc
S1^A^/HE^0^-Fc Lectin
B
1.0
0.8
)
m
n
(
g
n
i
d
n
i
B
0.6
μ
K
= 15.0 ± 2.2
M
D
0.4
0.2
0.0
0
10
20
30
Conc. (μM)
1.0
29.7 µM
0.8
)
m
n
(
g
n
i
d
n
i
B
14.8 µM
0.6
7.4 µM
0.4
3.0 µM
0.2
1.5 µM
0.0
0
50
100
150
200
Time (s)
sFig. 6. S protein of OC43-USA/1967 differs from BCoV-Mebus S in receptor-binding affinity and ligand preference. (A) Binding of BCoV and OC43 S1^A^-Fc fusion proteins to BSM was compared by sp-LBA as in Fig. 1. BSM is exceptionally rich in a variety of mono- and di-O-acetylated, 5-N-acetylated or -glycolylated Sias that can be selectively depleted by sialate-O-acetylesterase treatment with torovirus and coronavirus HE^+^-Fc proteins^6^. To assess potential differences in ligand preference between BCoV and OC43 S1^A^ binding was tested using BSM preparations, prepared and characterized previously^6^, that were depleted for mono-O-acetylated Sia species with porcine torovirus P4 HE to contain 7,9-di-O-Ac-Sias exclusively (7,9-di-O-Ac-BSM) or for 7,9-di-O-Ac-Sias with bovine coronavirus HE to contain 9-mono-O-Ac-Sias only (9-O-Ac-BSM). Virolectins, specific 9-O-Ac, 5-N-Ac-Sia (porcine torovirus PToV P4 HE^0^-Fc) or preferentially binding to 7,9-di-O-Ac-Sias (BCoV HE^0^-Fc) were included as controls. Note that BCoV S1^A^ binds preferentially to 7,9-di-O-Ac-Sia and that depletion of 9-mono-O-Ac-Sia by PToV P4 HE affects binding only modestly. Also note that binding of OC43 is strongly reduced by depletion of either type of O-Ac-Sia. (B) Binding affinities (K_D_) of BCoV measured by biolayer interferometry (BLI) with α2,6-linked 9-mono-O-Ac-SiaLacNAc. To allow monovalent 1 on 1 binding, S1^A^ domains were produced and purified by thrombin cleavage of S1^A^-Fc proteins at an engineered cleavage site. BLI was performed and binding affinities determined as described^3^. Note that although BCoV S1^A^ preferentially binds to 7,9-di-O-Ac-Sia, its K_D_ for α2,6-linked 9-mono-O-Ac-Sia such as are present in BSM^6^ is still 3-fold lower than that of OC43 S1^A^ ^3^.

### Slide 7
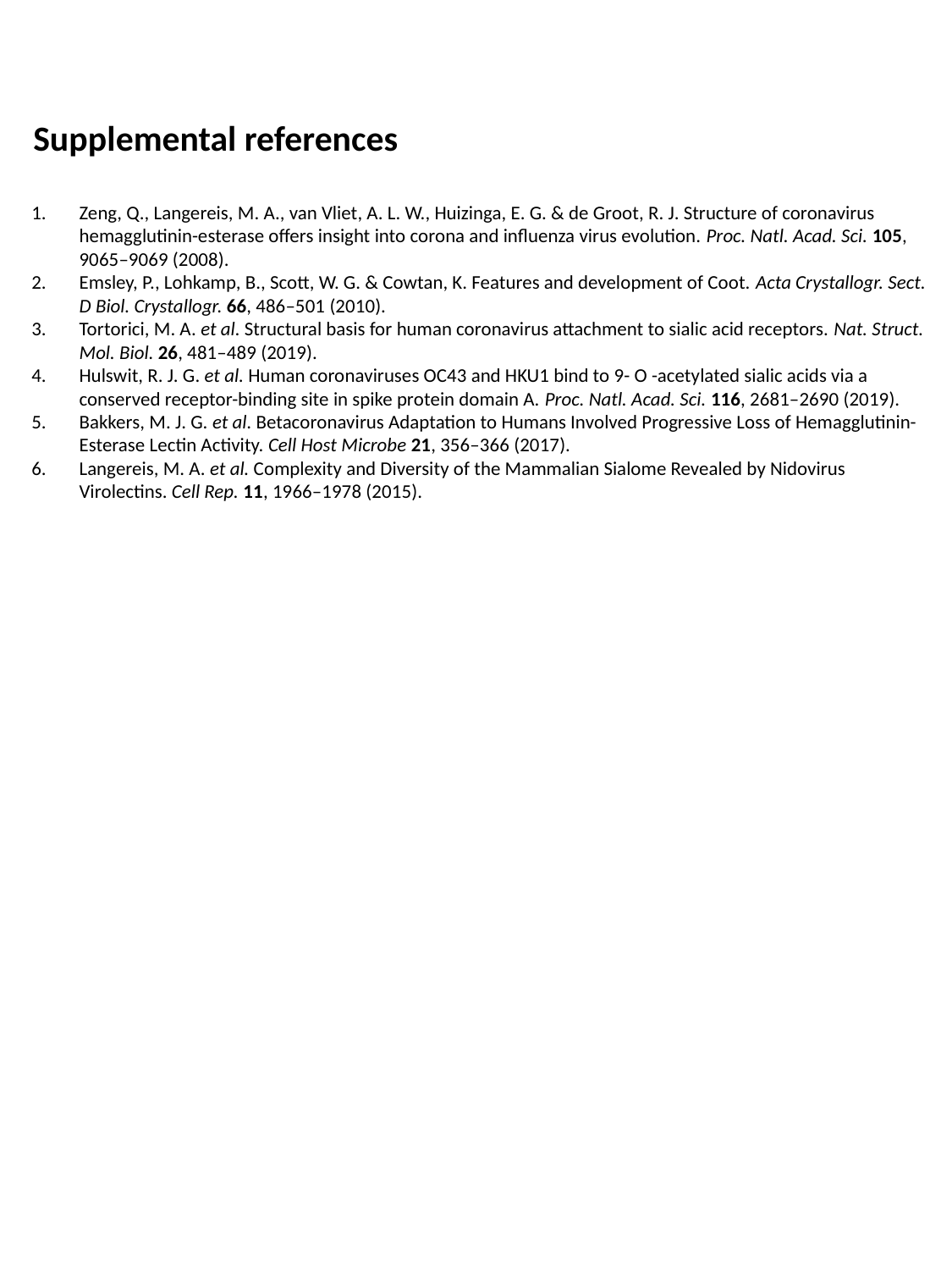
